# Post-translational modification patterns on β-myosin heavy chain are altered in ischemic and non-ischemic human hearts

**DOI:** 10.1101/2021.11.21.469462

**Authors:** Maicon Landim-Vieira, Matthew C. Childers, Amanda L. Wacker, Michelle Rodriguez Garcia, Huan He, Rakesh Singh, Elizabeth A. Brundage, Jamie R. Johnston, Bryan A. Whitson, P. Bryant Chase, Paul M.L. Janssen, Michael Regnier, Brandon J. Biesiadecki, J. Renato Pinto, Michelle S. Parvatiyar

**Author notes:** Co-equal first authors (Maicon Landim-Vieira, Matthew C. Childers). Correspondence should be addressed to: Dr. Michelle S. Parvatiyar, Ph.D., Florida State University College of Human Sciences 107 Chieftan Way, Tallahassee, FL 32306-1490.

## Abstract

Phosphorylation and acetylation of sarcomeric proteins are important for fine-tuning myocardial contractility. Here, we used bottom-up proteomics and label-free quantification to identify novel post-translational modifications (PTMs) on beta-myosin heavy chain (*β*-MHC) in normal and failing human heart tissues. We report six acetylated lysines and two phosphorylated residues: K34-Ac, K58-Ac, S210-P, K213-Ac, T215-P, K429-Ac, K951-Ac, and K1195-Ac. K951-Ac was significantly reduced in both ischemic and non-ischemic failing hearts compared to non-diseased hearts. Molecular dynamics simulations show that K951-Ac may impact stability of thick filament tail interactions and ultimately myosin head positioning. K58-Ac altered the solvent exposed SH3 domain surface – known for protein-protein interactions – but did not appreciably change motor domain conformation or dynamics under conditions studied. Together, K213-Ac/T215-P altered loop 1’s structure and dynamics – known to regulate ADP-release, ATPase activity, and sliding velocity. Our study suggests that β-MHC acetylation levels may be influenced more by the PTM location than the type of heart disease since less protected acetylation sites are reduced in both heart failure groups. Additionally, these PTMs have potential to modulate interactions between β-MHC and other regulatory sarcomeric proteins, ADP-release rate of myosin, flexibility of the S2 region, and cardiac myofilament contractility in normal and heart failure hearts.

## Introduction

The sarcomere is the smallest functional unit in striated muscle. The cardiac sarcomere is composed of thick and thin filament proteins that work together to generate force and shorten the sarcomere, and to regulate sarcomere contraction and relaxation in a Ca^2+^-dependent manner (1). The cardiac thick filament is composed of myosin II polymers accompanied by associated proteins, myosin-binding protein C (MyBP-C), titin, and obscurin (1–3). The major proteins of the cardiac thin filament include F-actin – containing myosin-binding sites – tropomyosin (Tm), and the cardiac troponin complex (cTn) (1). Myosin molecules of the thick filaments are constituted of six non-covalently associated polypeptides: two heavy chains and four light chains. The C-terminus of the two myosin heavy chains (MHCs) form an α-helical coiled-coil tail that extends toward the center of the thick filament backbone. The paired, N-terminal heads of the two MHCs are positioned at the surface – facilitating interactions with actin filaments. Between the heads and the tail is a coiled-coil that makes up the myosin S2 rod segment. The neck region of myosin is the site of accessory protein binding that consists of two pairs of light chains (LCs): essential (ELC) and regulatory (RLC). Near the N-terminus, the individual MHC forms a distinct globular structure – myosin S1 fragment – that interacts with the actin filament in a cyclic fashion (4). Myosin binds to actin and ATP and undergoes several conformational changes that are essential to its function: 1) ATP binding to myosin results in weak actin affinity and formation of the pre-power stroke state; 2) ATP hydrolysis occurs when myosin is dissociated from actin - while in a weak actin-binding state; and 3) binding of myosin to actin with hydrolyzed products (Pi and ADP) leads to the power stroke state (force generation) (5).

While myofilament proteins have distinct and key roles in cardiac sarcomere function, further regulation can be gained by post-translational modifications (PTMs). PTMs have been shown to alter the canonical structure, function, localization, and half-life of modified sarcomeric proteins (6). These modifications can instigate downstream effects on the functional properties of the myocardium thus providing a rapid, efficient, and energetically favorable mechanism to alter contractile function compared to isoform switching. The context-dependence of a modification may also be important, as it may be influenced by other PTMs on the same protein or other proteins. Furthermore, PTMs on sarcomeric proteins may be inert under normal conditions, but their functional importance can become evident alongside pathological conditions. Therefore, these normally “silent” PTMs represent novel targets for therapeutic intervention (7). Localized, spatially confined pools of kinases, acetyltransferases, and other protein modifiers have been identified as essential for the efficient modification of myofilament proteins. Both histone acetyltransferase (HAT; p300/CBP-associated factor PCAF) and histone deacetylase 4 (HDAC4) have been found localized at the sarcomeric matrix (8, 9). Furthermore, HDAC6 has also been shown to assume a sarcomeric localization (10). Of the PTMs identified on sarcomeric proteins, phosphorylation has been the most extensively characterized. However, acetylation, methylation, oxidation, SUMOylation, and ubiquitination have been reported as well – extending the repertoire of potential modifiers of the sarcomere (11, 12).

A number of sarcomeric proteins are phosphorylated including MyBP-C, troponin T (TnT), and troponin I (TnI) by cAMP-dependent protein kinase (PKA), and myosin regulatory light chain (RLC) by myosin light-chain kinase (MLCK) (7,13–15). Several kinases are implicated in phosphorylation of striated muscle Tm including tropomyosin kinase, PKA, and protein kinase C*ζ* (16–20). Remarkably absent from the list of phosphorylated sarcomeric proteins in human hearts is β-MHC, encoded by *MYH7*, the predominant isoform of myosin in the adult human heart. The massive size of β-MHC, ∼223 kDa, imposes limitations with current technological approaches and challenges our ability to obtain complete sequence coverage. In a study by Kawai et al. examining PTMs on *MYH6*, the predominant murine myosin isoform, 31 phosphorylation sites were identified in control hearts with a number of these residues not phosphorylated in HCM hearts (21). In addition, Jin et al. identified acetylated, methylated, and trimethylated residues in human β-MHC using size-exclusion chromatography (SEC)/middle-down mass spectrometry (MS) (22). Overall, detection of low-abundance PTMs on large proteins has remained elusive and protein enrichment strategies before Liquid Chromatography-Mass Spectometry (LC/MS) analysis or even greater instrument sensitivity can increase signal intensity of modified proteins (6, 23).

While many earlier studies were conducted in animals of varying species, examining changes in PTMs on sarcomeric proteins in the context of human heart disease may provide clues toward potential interventions. Identification of PTMs on *β*-MHC is the first step in discovering whether these modifications provide beneficial or adverse impacts on cardiac function and disease progression. With mass-spectrometry techniques increasing in sensitivity, we are becoming more successful at uncovering even low-abundance PTMs that may have significant impact on cardiac outcomes. As new reports emerge documenting the presence of PTMs in human sarcomeric proteins, the ensuing challenge remains: testing their *in vivo* functional consequences. Given the emerging importance of understanding the potential role of PTMs on cardiac muscle performance, we used LC/MS and molecular dynamics (MD) simulations to investigate novel PTMs sites in key functional regions of β-MHC in human hearts in healthy and diseased states. The present work provides insight into the potential of these modifications to fine tune the cardiac myofilament performance.

## Methods

### Human Heart Samples

Explanted donor human heart tissues were obtained from the Ohio State University Tissue program. The de-identified samples were obtained from patients that were 41-69 years of age with ischemic heart failure (I-HF) or non-ischemic heart failure (NI-HF), and healthy donors after informed consent. Clinical information about patient hearts that were utilized in this study is included in Supplemental Table 1.

### Heart Sample Preparation

Human heart tissue was homogenized in Laemmli buffer and separated by SDS PAGE (12%), stained with Coomassie blue, and bands that corresponded to β-MHC (∼223 kD) were excised (Supplemental Figure 1). The samples were homogenized in 1X Laemmli sample buffer with protease inhibitor cocktail, phosphatase inhibitor cocktail, 1 µM trichostatin A, 1 µM quisinostat, 5 mM nicotinamide, and 1 mM sodium vanadate.

### Mass Spectroscopy

Sample preparation - In-gel digests were performed for each excised sample using the ProteoExtract All-in-One Trypsin Digestion Kit (Cat. No. 650212; Calbiochem, EMD Millipore, Billerica, MA) according to manufacturer’s instructions. Briefly, excised gel pieces were de-stained in wash buffer, and dried at 90℃ for 15 min. Gel pieces were rehydrated in trypsin digestion buffer, and treated with a reducing agent for 10 min at 37℃. Samples were cooled to room temperature and then incubated in blocking reagent for 10 min at room temperature. Trypsin was added to a final concentration of 8 ng/µl and incubated for 2 h at 37℃ on an orbital shaker. Peptides were eluted in 50 µl 0.1% formic acid.

### Liquid chromatography-mass spectrometry (LC-MS)

Sample peptides were processed using an externally calibrated high-resolution electrospray tandem mass spectrometer (Thermo Q Exactive HF; Thermo Fisher Scientific, Waltham, MA) in conjunction with a Dionex UltiMate 3000 RSLCnano System (Thermo Fisher Scientific). The samples were masked during data acquisition. To load 5 µl of sample peptide was aspirated into a 50 µl loop and then loaded onto the trap column (Acclaim PepMap100 C18, 5 μm, 100 Å, 300 μm i.d. × 5 mm Cat #160454, Thermo Fisher Scientific). Separation on an analytical column (Acclaim PepMap RSLC 75 μm, and 15 cm nanoViper; Thermo Fisher Scientific) was conducted with a flow rate of 300 nL/min. A 60 min linear gradient from 3% to 45% B (0.1% formic acid in acetonitrile) was performed. The LC eluent was directly nanosprayed into Q Exactive HF mass spectrometer (Thermo Scientific). During the chromatographic separation, the Q Exactive HF was operated in a data-dependent mode and under direct control of the Thermo Excalibur 3.1.66 (Thermo Scientific). The MS data were acquired using the following parameters: 20 data-dependent collisional-induced-dissociation (CID) MS/MS scans per full scan (350 to 1700 m/z) at 60000 resolution. MS2 were acquired in centroid mode at 15000 resolution. Ions with single charge or charges more than 7 as well as unassigned charge were excluded. Raw data were searched with Proteome Discoverer 2.2 (Thermo Fisher Scientific) using Sequest HT and Mascot search engines and percolator as the PSM validator with species specific FASTA database. Phosphorylation and acetylation were used as a dynamic modification in SequestHT and Mascot and PTMs were scored by ptmRS node in Proteome Discoverer 2.2. MS1 based quantification of peptides was performed using Skyline 4.0. For access to the complete data set please utilize the following link: DOI (doi:10.5061/dryad.s4mw6m97g).

### Proteomics Data Analysis

For each peptide sequence, the area under the peak curve was measured in Skyline 20.1, an open source Windows client application software for targeted proteomics data analysis and bioinformatics (24). To determine the modification levels of each PTM site, the peak area of each modified peptide was measured and divided by the peak area of the unmodified common internal reference peptide (IRP) (Supplemental Figure 2). The ratio of the modified peptide to IRP was compared among human heart samples from non-diseased controls and patients with ischemic heart failure and non-ischemic heart failure. The IRP was chosen based upon recommendations from Sherrod et al. utilizing a known unmodified peptide between 7-20 amino acids (no methionine and cysteine residues), eluted across the chromatogram, and demonstrating consistent signal stability (25).

### Model Building

Starting coordinates for the wild type (WT) *β*-MHC motor domain simulations were obtained from an X-ray crystallography structure of the post-rigor, ATP state of *β*-MHC in the Protein Data Bank (PDB, www.rcsb.org) (PDB ID: 4DB1, 2.6 Å, residues 1-777) (26). Starting coordinates for the WT S2 fragment simulations were obtained from an X-ray crystallography structure of S2D (PDB ID: 2FXM, 2.7 Å, residues 838-963) (27). Missing heavy atoms were built using *Modeller* (28) and conformer ‘A’ was chosen among residues with multiple conformations in the PDB entries. Hg atoms were removed from the 2FXM structure and ANP•Mn was replaced with ATP•Mg in 4DB1. Starting coordinates for the post-translationally modified variants were obtained via *in silico* modification of the WT structures using the *leap* module of *AMBER*. There were two modified variants of 4DB1: 4DB1-K58-Ac (corresponding to acetylation of Lys 58) and 4DB1-K213-Ac/T215-P (corresponding to simultaneous acetylation of Lys 213 and phosphorylation of Thr 215). There was one modified variant of 2FXM: 2FXM-K951-Ac (corresponding to acetylation of Lys 951). For 2FXM, K951 residues in both chain A and B were modified.

### Force Field and Explicit Solvent Molecular Mechanics

All 4DB1 simulations were performed with the AMBER20 package (4, 5) and the ff14SB force field (29). Water molecules were treated with the TIP3P force field (7). Metal ions were modeled using the Li and Merz parameter set (30–32). ATP molecules were treated with parameters from Meagher *et al.* (33). Parameters for phosphothreonine (called ‘TPO’) and acetyllysine (called ‘ALY’) were obtained from Raguette *et al.* and Belfon *et al.,* respectively. The SHAKE algorithm was used to constrain the motion of hydrogen-containing bonds (34). Long-range electrostatic interactions were calculated using the particle mesh Ewald (PME) method (35).

### Pre-production protocols

Hydrogen atoms were modeled onto the initial structure using the *leap* module of *AMBER* and each protein was solvated with explicit water molecules in a periodic, truncated octahedral box that extended 10 Å beyond any protein atom. Na^+^ and Cl^-^ counterions were added to neutralize the systems and then 120 mM Na^+^ and Cl^-^ ions were added. Each system was minimized in three stages. First, hydrogen atoms were minimized for 1000 steps in the presence of 100 kcal mol^-1^ restraints on all heavy atoms. Second, all solvent atoms were minimized for 1000 steps in the presence of 25 kcal mol^-1^ restraints on all protein atoms. Third, all atoms were minimized for 8000 steps in the presence of 25 kcal mol^-1^ restraints on all backbone heavy atoms (N, O, C, and C atoms). After minimization, systems were heated to 310°K during 3 successive stages. In each stage, the system temperature is increased by ∼100°K over 100 ps (50,000 steps) using the NVT (constant number of particles, volume, and temperature) ensemble. During all heating stages, 25 kcal mol^-1^ restraints were present on the backbone heavy atoms (N, O, C, and C atoms). After the system temperatures reached 310°K, the systems were equilibrated over 5 successive stages using the NPT (constant number of particles, pressure, and temperature) ensemble. During each stage, the systems were equilibrated for 5.4 ns in the presence of restraints on backbone atoms. The strength of the restraints was decreased from 25 kcal mol^-1^during the first stage to 1 kcal mol^-1^ during the fourth stage. During the final equilibration stage, the systems were equilibrated in the absence of restraints.

### Molecular Dynamics Protocol

Production dynamics for conventional molecular dynamics (MD) simulations were then performed using the canonical NVT ensemble with an 8 Å nonbonded cutoff, and 2 fs time step. Coordinates were saved every picosecond. Simulations were run in triplicate for 500 ns each. Unless specified otherwise, simulations were analyzed separately, and the results of replicate simulations were averaged together. To account for potential equilibration effects, the first 100 ns were excluded from subsequent analyses.

### Implicit Solvent Simulations

2FXM contains a linear fragment of S2 spanning ∼130 residues. Due to the length of this fragment, we performed implicit solvent simulations of 2FMX using the Generalized Born model. This approach was previously used to study S2 fragments and our methods were chosen to best match earlier simulations (36). Simulations were performed using the GB model described by Mongan *et al.* (37).

### MD Analysis

The C*_α_* RMSD, C*_α_* RMSF, SASA, interatomic distances, and interatomic contacts were calculated with *cpptraj* (38). The C*_α_* RMSD was calculated after alignment of all The C*_α_* atoms to the minimized structure. The C*_α_* RMSF was calculated about average MD structures for each simulation. Two residues were considered in contact with one another if at least one pair of heavy atoms were within 5 Å of one another. All protein images were prepared using UCSF Chimera (39, 40). Electrostatic potentials of molecular surfaces were generated using default parameters for the *Couloumbic Surface coloring* method in *UCSF Chimera*.

### Statistical Analysis

Mass spectrometry and MD simulations data analyses were performed using one-way ANOVA, followed by Bonferroni post hoc test and Student’s *t*-test, respectively. The limits of exclusion for this study is that all data obtained would be reported even if outlier values exceed +/-2 standard deviation.

## Results

### Human Heart Data Bank

The patient groups from whom explanted heart tissues were collected included explanted hearts from healthy donors (non-failing, NF) and end-stage heart failure patients, who were reported to have either ischemic heart failure (I-HF) or non-ischemic heart failure (NI-HF). The demographics for the patients included information on age, gender, and race. Patients’ ages ranged from 41-69 years with each condition represented (NF, NI-HF and I-HF) (Supplemental Table 1), and were similarly distributed among the three groups. In an attempt to provide gender and race balance, one female was included in each group and at least one African American. We utilized mass spectrometry to investigate whether PTMs could be found on β-MHC isolated from these hearts and utilized MD simulations to better understand functional impacts of these PTMs that may have in human cardiac disease presentation. For work flow, see the schematic in Figure 1.

**Figure 1.**
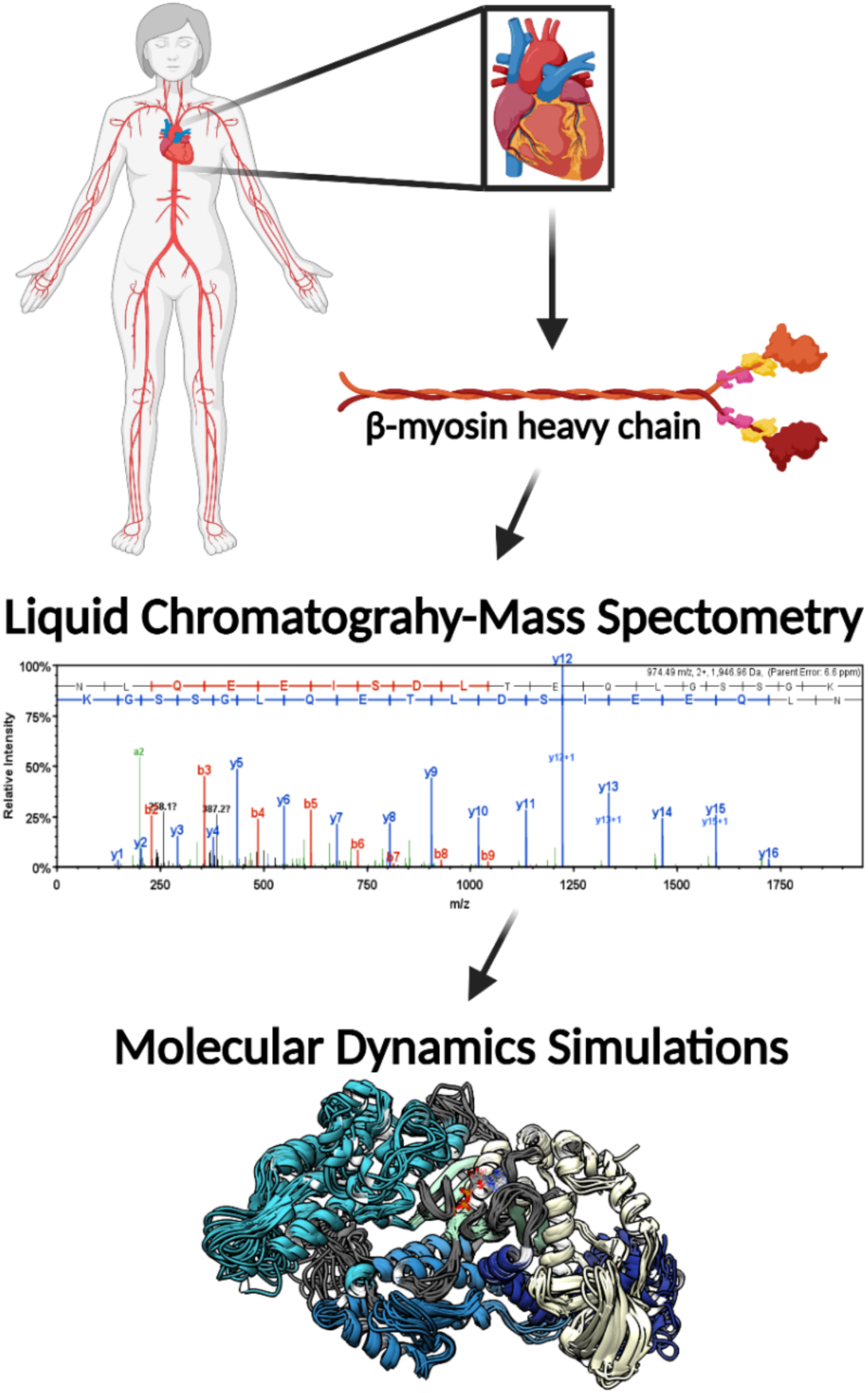
Illustrative schematic of the integrative approaches used to identify novel PTMs on human β-MHC and investigate their roles on cardiac muscle regulation. De-identified human heart samples were obtained from non-failing, ischemic heart failure, and non-ischemic heart failure patients. The presence of PTMs on human β-MHC was confirmed by LC/MS. b-ions and y-ions indicate N-terminal and C-terminal ions, respectively. MD simulations were used to further understand the functional significance of the newly-identified PTMs on β-MHC. The schematic was generated using BioRender.com.

### Identification and Location of PTMs on human cardiac β-MHC

Unique high confidence peptides bearing PTMs were identified using bottom-up mass spectrometry data from non-failing, ischemic, and non-ischemic failing human hearts (Figures 2 and 3, and Supplemental Figures 3 and 4). The samples were purified using SDS-PAGE and were digested using in-gel trypsinization (Supplemental Figure 1). We report eight PTMs: six acetylated lysines and two phosphorylated residues, one serine and one threonine: K34-Ac, K58-Ac, S210-P, K213-Ac, T215-P, K429-Ac, K951-Ac, and K1195-Ac (Tables 1 and 2). Six of these PTMs are distributed throughout the myosin motor domain (Figure 4A) and two within the tail (Figure 4B). The myosin motor is comprised of four domains: the N-terminal domain, the upper and lower 50 kDa domains, and the converter arranged around a central *β*-sheet (Figure 4A). The structure of myosin gives rise to three functional regions. The actin binding cleft is formed between the upper and lower 50 kDa domains and interacts with the thin filament. The nucleotide binding pocket formed between the upper 50 kDa domain and binds actin filaments. Several loops near or within the nucleotide binding pocket coordinate the nucleotide and transmit structural information throughout the structure. These include the phosphate binding loop (p-loop), which coordinates the terminal phosphates of ATP/ADP; switch 1 and switch 2, which communicate with the upper and lower 50 kDa domains, respectively; and loop 1, which regulates ATP cycling. The converter-tail junction formed by the converter domain, tail, and N-terminal domain converts small-amplitude structural changes in the motor domain into large-amplitude lever arm motion. A schematic of the *β*-MHC protein is included in Supplemental Figure 3 indicating key regions of the protein along with the newly identified PTMs. In Figure 4A, the three-dimensional structure of the human *β*-MHC sequence complexed with Mn-AMPPNP (PDB 4DB1) was used to model the PTM sites at K58-Ac and doubly-modified peptide K213-Ac/T215-P. This structure represents a post-rigor ATP-bound state of myosin and was used to assess proximity of the modified residues to structurally important regions in the myosin motor domain. An S2 fragment structure (PDB: 2FXM) was used to model K951-Ac.

**Figure 2.**
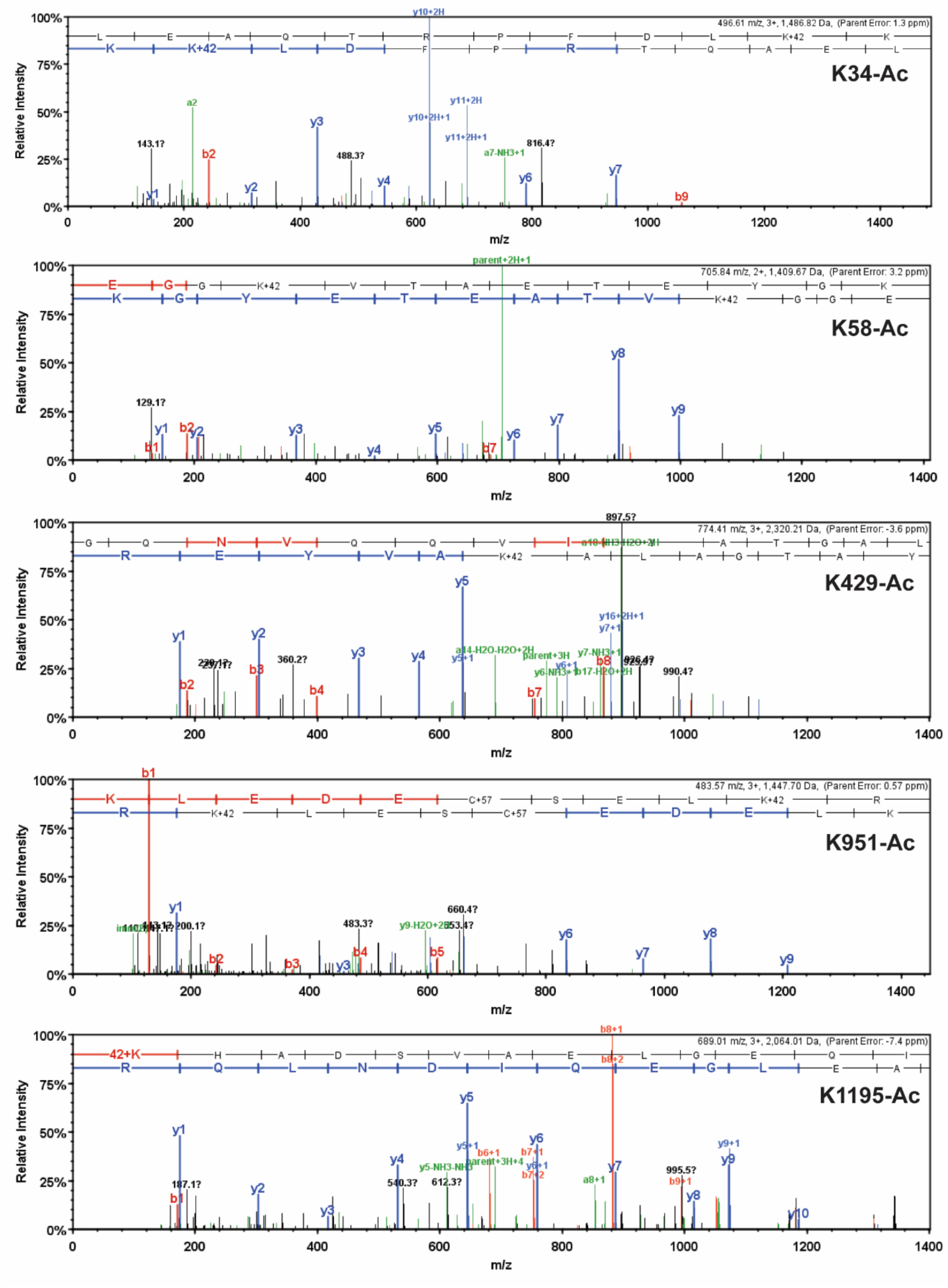
Detailed MS/MS spectrums of acetylated human β-MHC peptide sequences. MS/MS spectrums of trypsin-digested, acetylated β-MHC peptide sequences 24-35 (K34-Ac, m/z 496.61), 55-67 (K58-Ac, m/z 705.84), 414-434 (K429-Ac, m/z 774.41), 942-952 (K951-Ac, m/z 483.57), and 1195-1212 (K1195-Ac, m/z 689.01). b-ions and y-ions indicate N-terminal and C-terminal ions, respectively.

**Figure 3.**
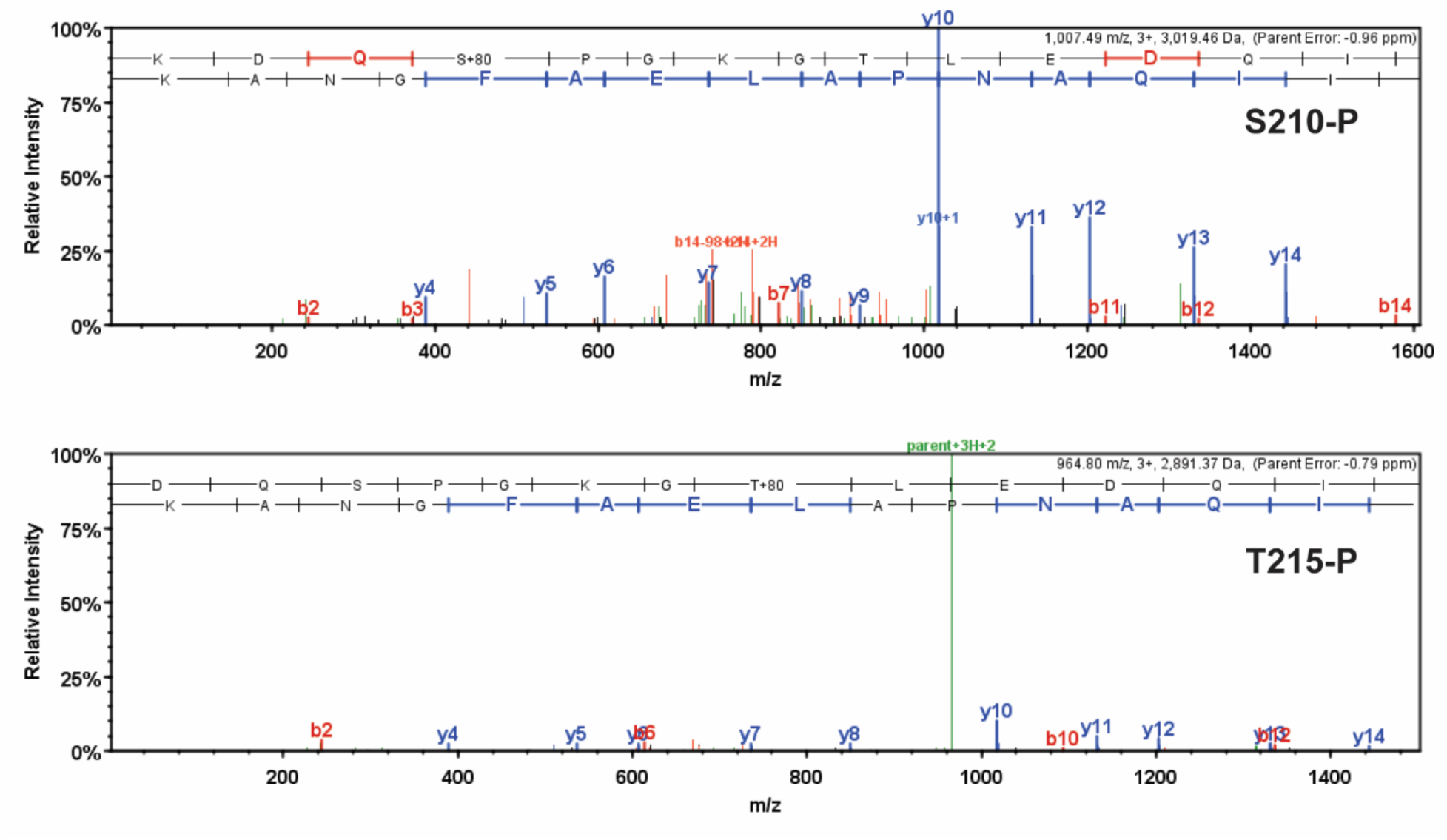
Detailed MS/MS spectrums of phosphorylated human β-MHC peptide sequences. MS/MS spectrums of trypsin-digested, phosphorylated β-MHC peptide sequences 207-234 (S210-P, m/z 1007.49) and 208-234 (T215-P, m/z 964.80). b-ions and y-ions indicate N-terminal and C-terminal ions, respectively.

**Figure 4.**
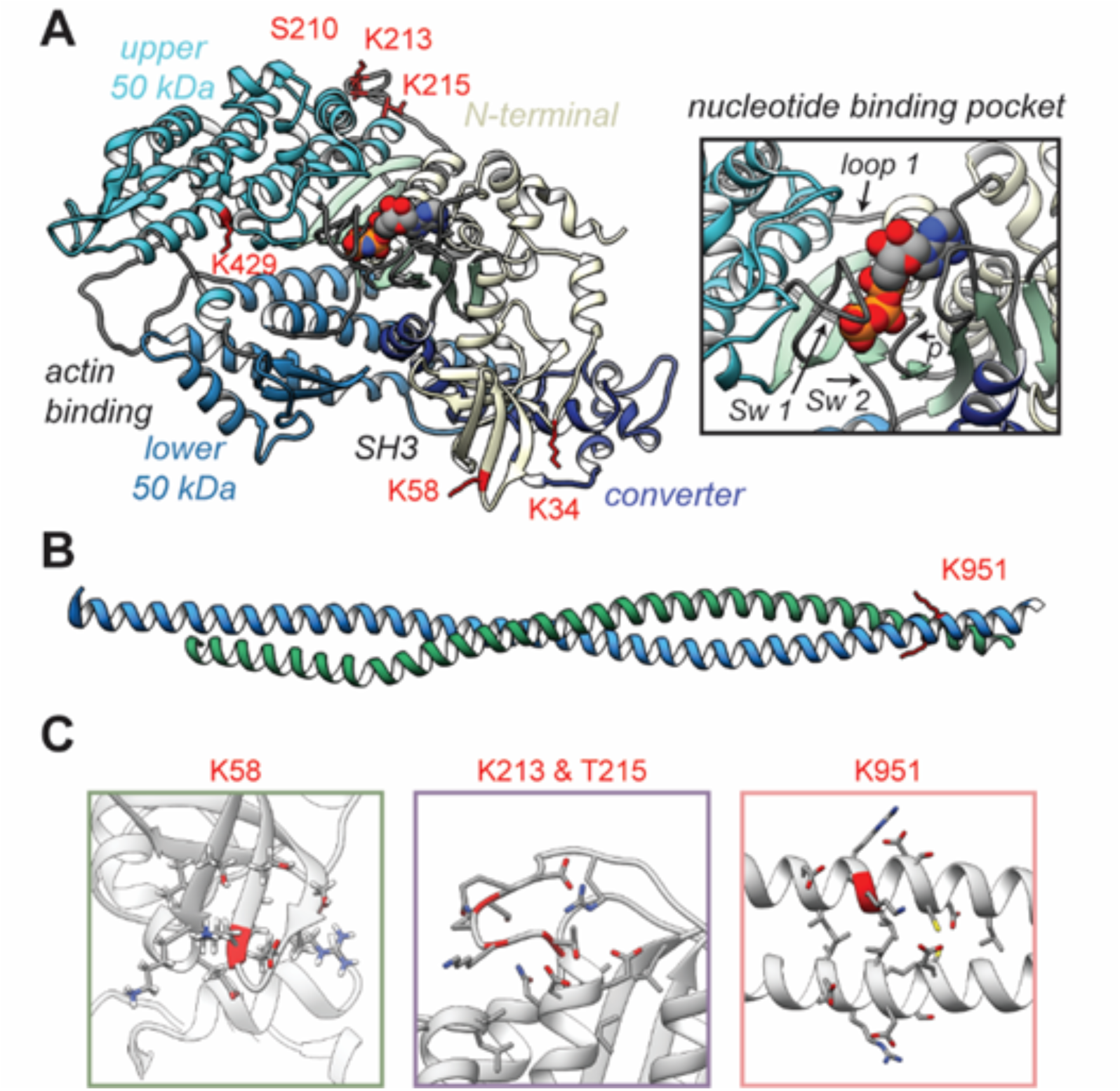
Structural models of PTMs on β-MHC. X-ray crystal structures of the *β*-MHC. In (A) a post-rigor X-ray structure was used to model K58-Ac, K213-Ac, T215-P PTMs. The four motor subdomains: the N-terminal domain (yellow), upper 50 kDa domain (cyan), lower 50 kDa domain (light blue), and converter domain (dark blue) and function sites they form are labeled. Inset highlights the nuceltide binding pocket and functional loops (loop 1, Switch 1, Switch 2, phosphate binding loop). B) An X-ray structure of an S2 fragment was used to model the K951-Ac PTM and served as the initial conformations of MD simulations. In A and B, residues with reported PTMs are shown and colored red. C) The colored boxes display side chain atoms in the vicinity of the modified (red ribbon) residues for K58 (green), K213/T215 (purple), and K951 (pink).

**Table 1.**
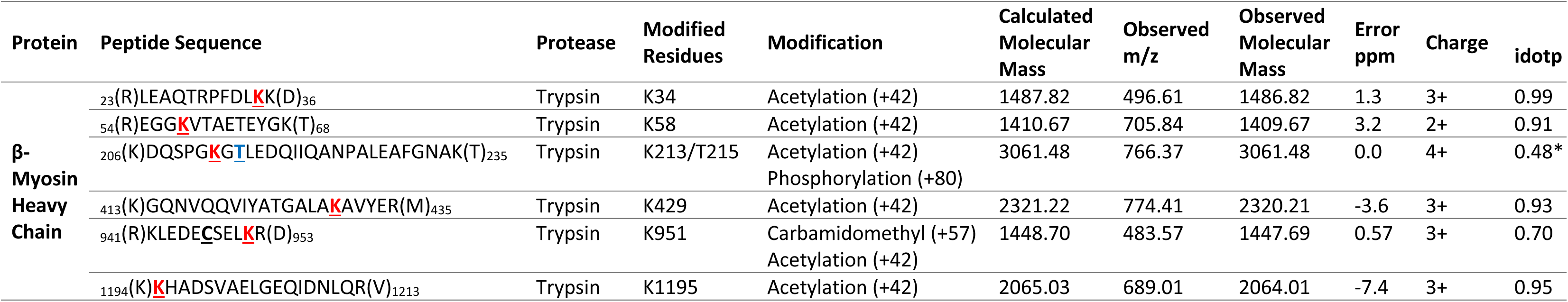
LC-MS/MS analysis of trypsin-digested human β-MHC sequences containing acetylated residues. An isotope dot product (idotp) value below 0.5 denotes a detectable but not quantifiable peptide sequence; **<u>C</u>** = carbamidomethyl. Acetylated residues are highlighted in red while phosphorylated residues are found in blue.

**Table 2.** LC-MS/MS analysis of trypsin-digested human β-MHC sequences containing phosphorylated residues. An isotope dot product (idotp) value below 0.5 denotes a detectable but not quantifiable peptide sequence; **<u>C</u>** = carbamidomethyl. Acetylated residues are highlighted in red while phosphorylated residues are found in blue.

| Protein | Peptide Sequence | Protease | Modified Residues | Modification | Calculated Molecular Mass | Observed m/z | Observed Molecular Mass | Error ppm | Charge | idotp |
| --- | --- | --- | --- | --- | --- | --- | --- | --- | --- | --- |
| **β-Myosin Heavy Chain** | <sup>206</sup>(K)KDQ**S**PGKGTLEDQIIQANPALEAFGNAK(T)<sub>235</sub> | Trypsin | S210 | Phosphorylation (+80) | 3020.47 | 1007.49 | 3019.46 | -0.96 | 3+ | 0.99 |
|  | <sup>206</sup>(K)KDQSPG**KGT**LEDQIIQANPALEAFGNAK(T)<sub>235</sub> | Trypsin | K213/<br>T215 | Acetylation (+42)<br>Phosphorylation (+80) | 3061.48 | 766.37 | 3061.48 | 0.0 | 4+ | 0.48<br>* |
|  | <sup>207</sup>(K)DQSPGKG**T**LEDQIIQANPALEAFGNAK(T)<sub>235</sub> | Trypsin | T215 | Phosphorylation (+80) | 2892.37 | 964.80 | 2891.37 | -0.79 | 3+ | 0.99 |

Residue K34 is located between the SH3-like domain and the converter domain. K58 is located within the SH3-like domain, which typically serves as a module for protein-protein interactions. Therefore, it is anticipated that acetylation of either K34 or K58 may interfere with normal interactions of this domain. Residues S210, K213 and T215 are sequentially close in the primary structure, and it is interesting to note that they lie together on one face of the myosin motor domain near the ATP binding pocket in a region called loop 1. This loop is notable for its influence on ATP/ADP cycling (41). It is plausible therefore, to expect that these modifications may influence ATP binding and/or Pi and ADP release dynamics. S210-P was found to exist as a single modification only. The location and the dynamics between modifications of residues K213-Ac and T215-P are interesting as they appear to be co-modifications. Although K213-Ac was only found coincident with phosphorylation of T215, T215-P was also found on its own, without K213-Ac. The K213-Ac/T215-P modified sequence (amino acids 207-234, m/z 766.36) was detectable as an isotope dot product (idotp) with a value below 0.5, and therefore was unquantifiable (Supplemental Figure 4). The acetylated residue K429 is located within the myosin head-like domain at the actin-binding interface. The remainder of the modified residues K951 and K1195 are located within the coiled-coil of the S2 region, although K1195 is missing from the model in Figure 4A. The panels within Figure 4B show a close up views of secondary structural elements and side chain interactions that neighbor the K58-Ac, K213-Ac, T215-P, and K951-Ac modifications.

### Normalized Peak Areas of the Post-Translational Modification Sites

To investigate the potential significance of these newly identified PTMs for cardiac function, we assessed their abundance in the human heart samples we analyzed. The relative abundance of the modifications was determined by calculating peak areas of modified peptide and normalizing to IRP peak area (amino acids 1504-1521, m/z 974.49). The MS/MS spectrum of the trypsin-digested common IRP is shown in Supplemental Figure 2 and b-ions indicate N-terminal fragment ions and y-ions indicate C-terminal fragment ions.

Overall, the ratios of modified PTM sites were variable, ranging between 1–14 in the non-failing (NF) donor hearts, with a tendency for decreased abundance in failing hearts. In Figure 5, the ratio of modified/IRP is shown for peptide 1 (K34-Ac), peptide 2 (K58-Ac), peptide 3 (K429- Ac), peptide 4 (K951-Ac) and peptide 5 (K1195-Ac). Beneath each histogram are the respective tryptic peptides where the reported PTMs were found. The residues shown in brackets are the trypsin digestion sites. Of interest is PTM K951-Ac as acetylation at this site is significantly decreased with ratios of approximately 14 in control hearts and 5 in failing hearts. Following the same trend (but not statistically significant) is K1195 also in the tail region. Both K34-Ac and K429-Ac are found in the myosin motor domain where they may be protected from HDAC activity. K429-Ac is a low abundance PTM with ratios of modified peptides/IRP ratios of approximately 1 and 1.4. This site is buried within the actin-binding cleft and may have been modified prior to protein folding. K34-Ac on the other hand is more abundant with approximate modified peptide ratios of 5 to 10. Perhaps the most interesting observation can be made for the co-modified peptide containing K58-Ac as it may be more susceptible to removal by HDACs or lower HAT activity under ischemic heart failure conditions.

**Figure 5.**
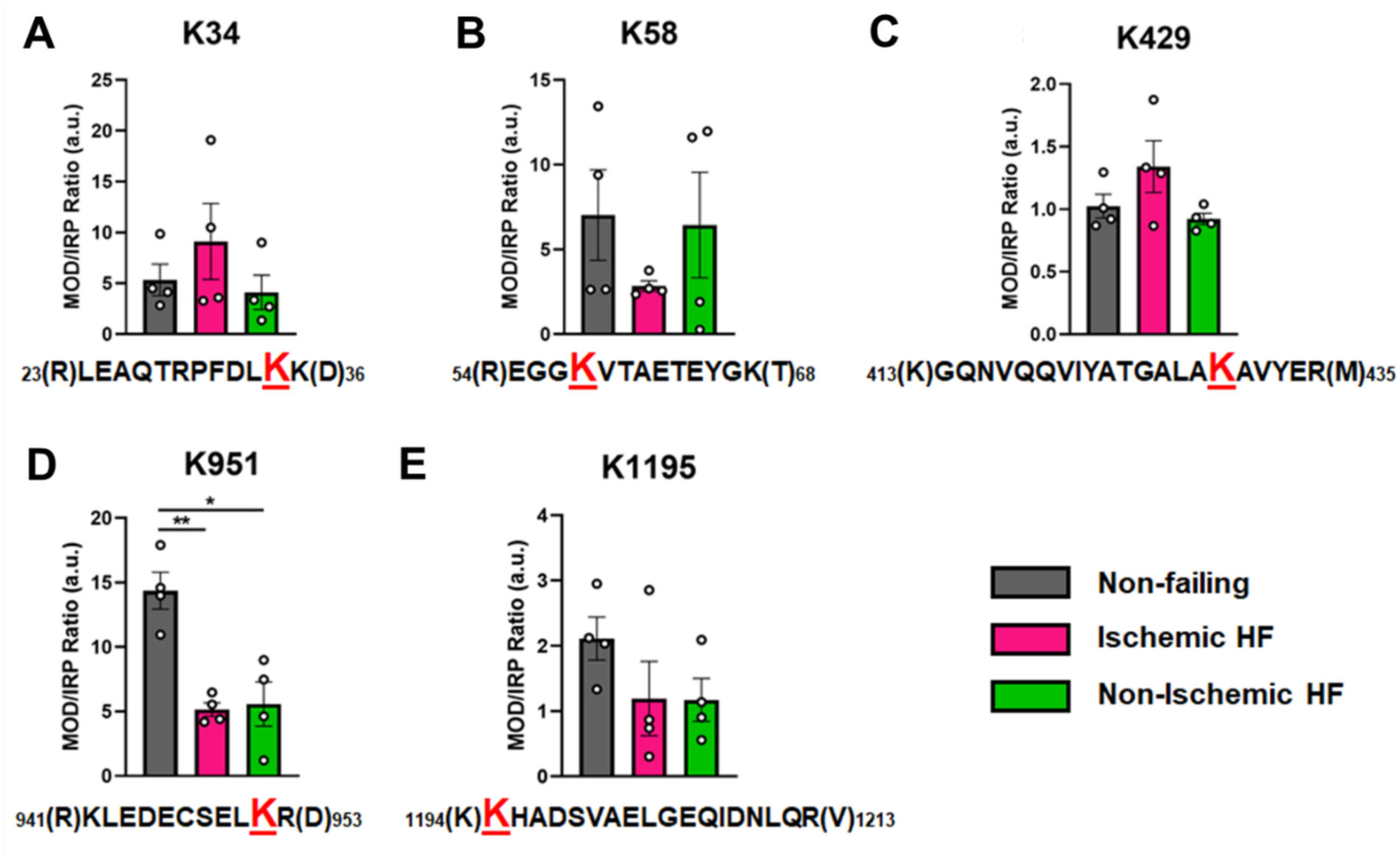
Calculations of PTM occupancy of acetylated residues on β-MHC. Peak areas of all modified peptide sequences (MOD) were normalized to the peak area of a common internal reference peptide sequence (IRP, 1504-1521). In (A) peptide 1 sequence with site of acetylated lysine residue K34 indicated in red. (B) peptide 2 shown with site of acetylated lysine residue K58 shown as red. In (C) peptide 3 sequence is shown with acetylated lysine residue K429. In (D) peptide 4 sequence with acetylated lysine residue K951 shown in red. In (E) peptide 5 sequence shown with acetylated lysine residue K1195 indicated in red. In the histograms different human heart samples are indicated with non-failing donor hearts (gray), ischemia-induced heart failure (pink) and non-ischemia induced heart failure (green). Trypsin-cutting sites are shown between parentheses. Data are expressed as mean ± SEM. Statistical analysis was performed by one-way ANOVA followed by Bonferroni post hoc test, * P < 0.05, n=4.

In Figure 6, the modified/IRP ratios are shown for the phosphorylated peptides. The average ratios of T215-P ranged from 3 to 2, while in the control non-diseased hearts the modification was nearly undetectable in two samples. The modifications on the other high confidence peptides may have functional significance, although the ratio of significantly modified/IRC was not altered in diseased (I-HF and NI-HF) compared to NF hearts (Figures 5 and 6). For this reason, we expect they may have a limited role in driving pathogenesis, but may still have functional relevance. Another phosphorylated residue was the phospho-serine at S210 on peptide 6, with ratios of approximately 1 to 2 modified peptide/IRP in all groups. T215 on peptide 7 with had ratios of approximately 1 to 2 of the modified peptides/IRP in all the groups (Figure 6). When examining the 3D structure of the β-MHC motor domain, we observed three modifications – at the lateral face of the myosin motor domain – with potential importance due to their proximity to the ATP binding pocket: 1) S210-P (single), 2) T215-P (single), and 3) K213-Ac plus T215-P (double). However, the question remains whether PTMs with low occupancy on a protein such as β-MHC, may in fact have subtle yet important roles in fine tuning of its function. Additional factors to consider are whether specific PTM sites reported here are more labile and prone to action by phosphatases and deacetylases, which may have increased activity in the failing heart.

**Figure 6.**
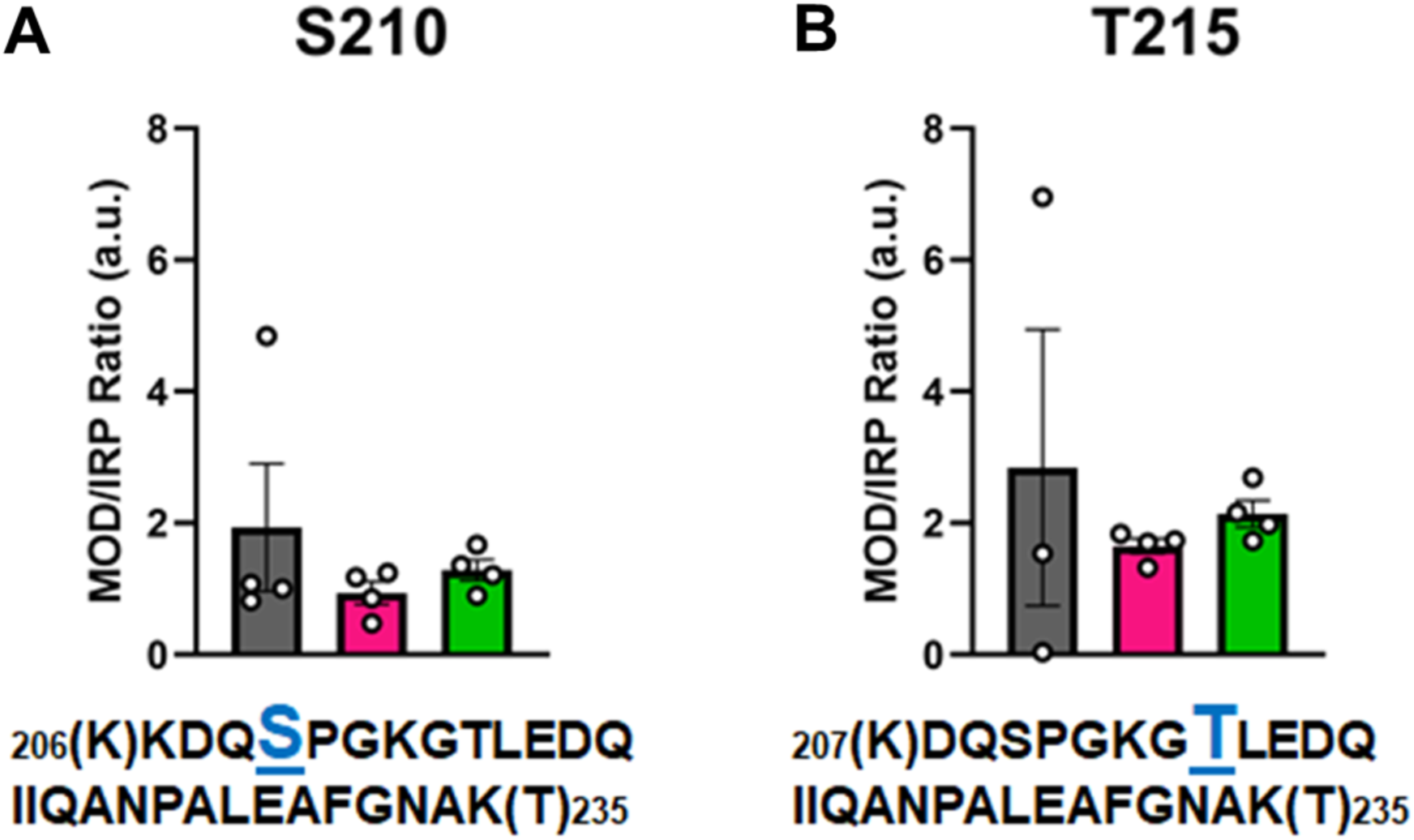
Calculations of PTM occupancy of phosphorylated residues on β-MHC. Peak areas of all modified peptide sequences (MOD) were normalized to the peak area of a common internal reference peptide sequence (IRP, 1504-1521). In (A) peptide 1 sequence shown with phosphorylated serine indicated in blue. In (B) peptide 2 sequence shown with phosphorylated threonine shown in blue. Trypsin-cutting sites are shown between parentheses. Data are expressed as mean ± SEM. Statistical analysis was performed by one-way ANOVA followed by Bonferroni post hoc test, n=3-4.

### Modeling the Potential Functional Changes in β-MHC due to Identified PTMs

Model building and MD were used to study putative relationships between PTMs of β-MHC and cardiovascular disease. Here, we performed MD simulations of unmodified and modified β-MHC (K58-Ac, K213-Ac/T215-P, K951-Ac) of PTMs that existed in functionally significant regions of the protein seen in Figure 4). For details regarding simulation runs refer to Tables 3 and 4. From Figure 4, it can be seen that phosphorylation and acetylation sites were distributed throughout the structure of the motor domain and S2 fragment.

**Table 3.** Inventory of β-MHC PTM simulations. Each row corresponds to a simulated system and reports the modifications that were made and the extent of MD sampling.

| **ID** | **PDB** | **Condition** | **Runs** | **Length per Run**<br>**(ns)** | **Net Sampling**<br>**(ns)** |
| --- | --- | --- | --- | --- | --- |
| 4DB1 Unmodified | 4DB1 | Unmodified | 3 | 500 | 1,500 |
| 4DB1 K58-Ac | 4DB1 | K58Ac | 3 | 500 | 1,500 |
| 4DB1 K213-Ac/T215-P | 4DB1 | K213Ac, T215P | 3 | 500 | 1,500 |
| 2FXM K951 | 2FXM | Unmodified | 3 | 500 | 1,500 |
| 2FXM K951-Ac | 2FXM | K951Ac | 3 | 500 | 1,500 |

#### Lysine 58

The overall motor domain in the K58-Ac simulations sampled conformations similar to those sampled by the WT simulations. The K58-Ac simulations had similar average C*_α_* to the unmodified simulations (Table 4). We did not detect any appreciable differences in overall conformational sampling between the WT and K58-Ac simulations. Next, we examined local changes in structure caused by K58-Ac. Lys 58 is located within the SH3-like domain of MYH7. SH3 domains are typically modules of protein-protein interaction and canonically bind to proline-rich sequence with polyproline helical structure (42). In typical SH3 domains, the poly-Pro binding site is formed by several small pockets intercalated between RT-loop and two β-strands and include conserved Tyr and Trp residues. These residues are conserved among many myosins, but not in the N-terminal SH3 domain of MYH7, where they are replaced by K58 and K72. Despite these substitutions, potential binding pockets remain on the surface of MYH7’s SH3 domain: one is formed between K58 and K72 and another formed between K72, T70, T60, S53, and E55 (Figure 7A and C). We note that there is no direct structural evidence that these residues do form poly-Pro peptide binding pockets and have instead inferred this structural role from homology to other SH3 domains. In the unmodified simulations both putative binding pockets remained intact and accessible to potential binding partners. In the K58-Ac simulations, however, the uncharged, acetylated Lys alternated between two conformations. The first was a ‘native-like’ conformation in which the side-chain projected into solvent and made transient hydrophobic interactions with the aliphatic portion of the neighboring K72 and a transient salt bridge with the neighboring E55 (Figure 7A). The second was an ‘intercalated’conformation in which the aliphatic portion of K58 was sandwiched in between K72 and T70 (Figure 7A and C). Formation of the ‘intercalated’ conformation became more likely as the acetylated Lys was capable of forming hydrogen bonds with K72 and T70 side chains and was no longer subject to electrostatic repulsion from K72. The ‘intercalated’ conformation was associated with a disruption of the native contact network among SH3 residues,an increase in the solvent accessible surface area (SASA) of residue K58 (partially attributable to the intrinsic increase in SASA of acetylated Lys), and a decrease in the SASA of K72 (Figure 7B). Importantly the ‘intercalated’ conformation abolished one of the putative poly- Pro binding pockets on the surface of SH3 (Figure 7C). This suggests that K58 acetylation would affect the ability of the SH3 domain to bind its targets.

**Figure 7.**
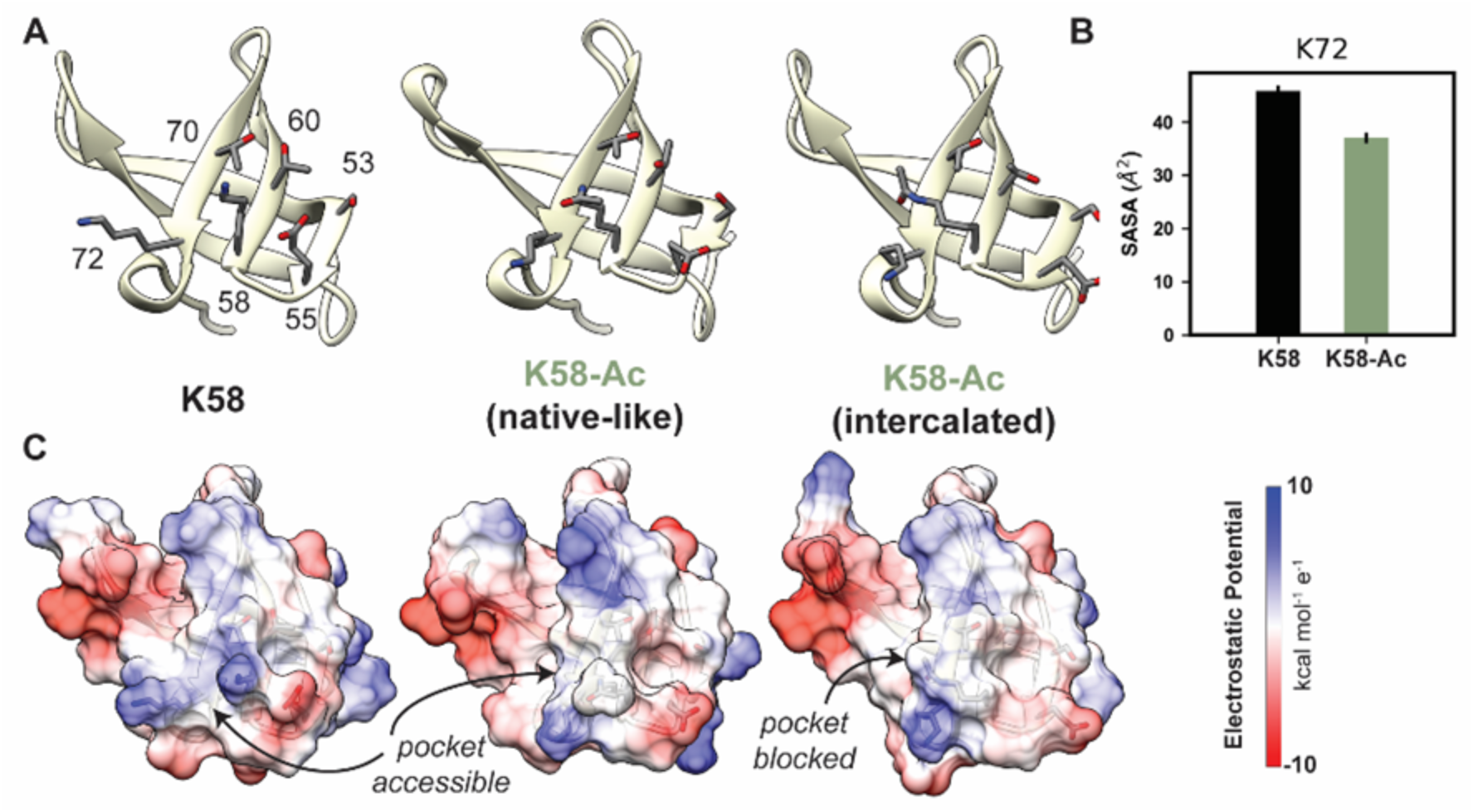
K58-Ac Altered the Solvent Exposed Surface of Myosin’s SH3-like Domain. (A) Representative structural changes in the SH3 domain associated with K58-Ac are shown in the endpoint structures of one unmodified (‘K58’) and two modified (‘K58-Ac’) simulations. K58- Ac formed increased interactions with T70, V71, and K72 in the neighboring strand and transiently formed an ‘intercalated’ conformation where the side chain was inserted between T70 and K72. (B) The transiently formed intercalated conformation led to a decrease in the solvent accessible surface of K72. (C) Modification of K58 also altered the electrostatic potential of the SH3 domain surface and in the intercalated conformation, K58-Ac blocks a surface pocket. The molecular surfaces in C correspond to the same structures shown in A and the electrostatic potential was calculated using *Chimera’s* Couloumbic surface coloring method. Statistical analysis was performed using Student’s t-test and n=3 simulations were run per modification.

**Table 4.** Average Cα RMSD Values. The C*α* RMSDs of each MD snapshot versus the reference structure were averaged together per simulation.

| ID | Run Number | C<sub>α</sub> RMSD (Å) |
| --- | --- | --- |
| 4DB1 Unmodified | 1 | 3.9 ± 0.3 |
|  | 2 | 3.2 ± 0.3 |
|  | 3 | 4.1 ± 0.3 |
| 4DB1 K58-Ac | 1 | 3.8 ± 0.3 |
|  | 2 | 3.6 ± 0.2 |
|  | 3 | 4.1 ± 0.5 |
| 4DB1 K213-Ac/T215-P | 1 | 3.5 ± 0.4 |
|  | 2 | 3.6 ± 0.4 |
|  | 3 | 3.6 ± 0.3 |
| 2FXM Unmodified | 1 | 6.9 ± 2.4 |
|  | 2 | 7.5 ± 2.3 |
|  | 3 | 7.2 ± 2.4 |
| 2FMX K951-Ac | 1 | 10.8 ± 2.8 |
|  | 2 | 9.2 ± 2.5 |
|  | 3 | 10.1 ± 2.0 |

#### Lysine 213/Threonine 215

Lys 213 and Thr 215 are both located in loop 1 of myosin S1, which is comprised of residues 199-215 in β-MHC (Figure 8). Loop 1 connects two α-helices that line the nucleotide-binding pocket: one in the N-terminal domain and one in the upper 50 kDa domain. The opposite end of the N-terminal helix forms a phosphate-binding loop that binds the phosphate groups in ATP/ADP. The opposite end of the upper 50 kDa helix forms switch 1, which also interacts with ATP/ADP and is also responsible for communicating structural information between the nucleotide-binding pocket and the actin-binding cleft. Thus, loop 1 is intimately connected with structure and dynamics of the nucleotide binding pocket. Loop 1 is flexible and is rarely resolved in X-ray structures. Residues 205-211 are not present in the 4DB1 X-ray crystal structure and thus were built into our structural model prior to performing our simulations. MD is unique in that it can reveal aspects of loop 1 that are ‘hidden’ crystallographically. The X-ray structure contains backbone coordinates for both K213 and T215, but only the C_b_ atom of K213 is resolved. In the X-ray crystal structure, K213 is oriented toward the upper 50 kDa domain and T215 is a helix-capping residue: the side chain of T215 interacts with the backbone NH group of D218 and the backbone of T215 interacts with the backbone of both D218 and Q219. In our model, the reconstructed side chain of K213 did not form additional interactions. In simulations without PTMs, T215 retained its role as a helix capping residue and the crystallographic interactions between T215, D218, and Q219 were preserved K213 formed a transient salt bridge with D337, transient hydrogen bonds with N334, and transient hydrophobic interactions with V338 (Figure 8A). These interactions tethered the C-terminal end of loop 1 to the upper 50 kDa domain. Residue-residue interactions were altered in the K213-Ac/T215-P simulationsK213 lost interactions with N334/D337 in the upper 50 kDa domain and instead formed interactions with T215. T215-P, now negatively charged, gained interactions with the postiviely charge R204 and K206. The altered amino acid interactions caused loop 1 to sample from a distinct conformational ensemble in the PTM simulations: it formed a more compact structure that involved more interactions with other loop 1 residues as opposed to an extended conformation with interactions to other regions of myosin.

**Figure 8.**
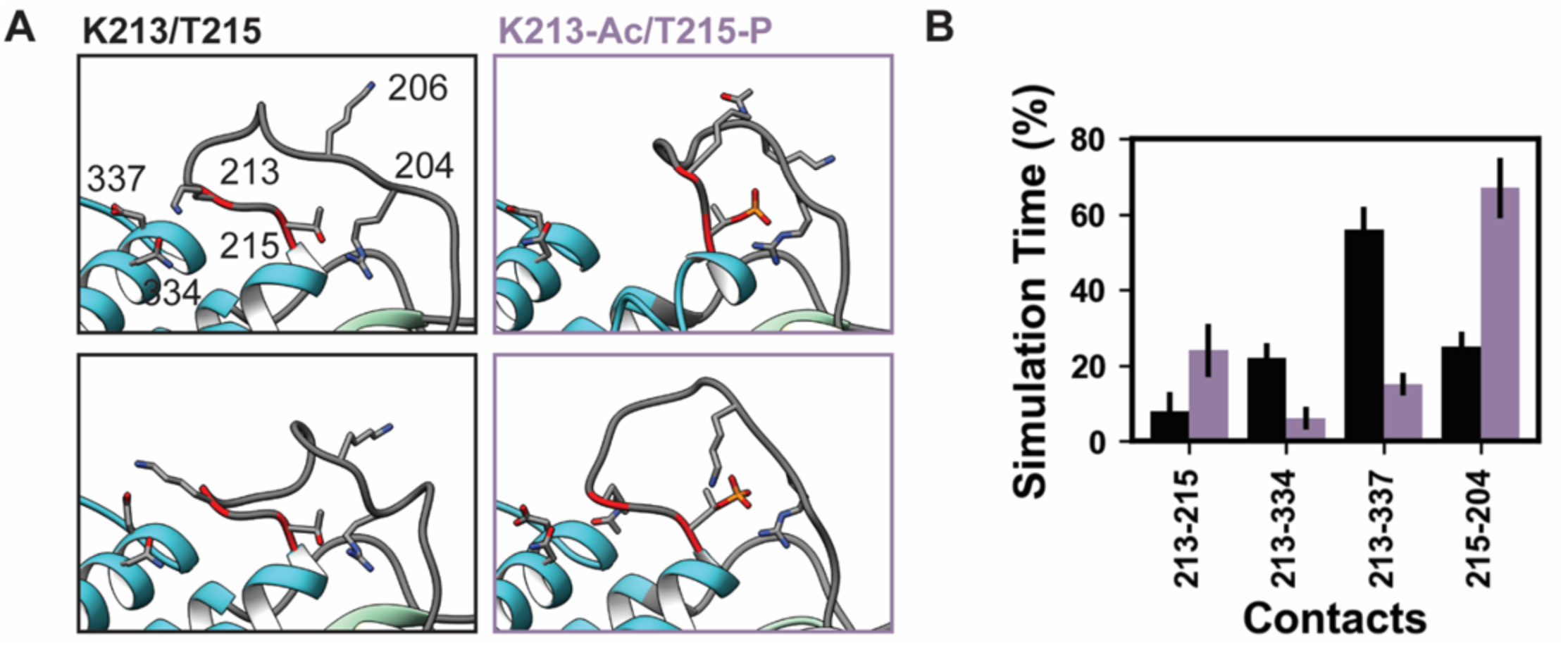
K58-Ac / T215-P Altered Loop 1 structure and dynamics . MD snapshots in (A) show representative structures from the unmodified (K213/T215) and modified (K213-Ac/T215-P) simulations. In the unmodified simulations, the loop makes long lasting interactions with the upper 50 kDa domain (teal) of myosin. In the modified simulations, became more compact and made fewer interactions with the upper 50 kDa domain. (B) The K213-Ac and T215-P PTMs altered the duration of inter-residue contacts made by K213 and T215: fewer interactions were made with the upper 50 kDa domain of myosin and more enduring interactions were made with other loop 1 residues. Statistical analysis was performed using Student’s t-test and n=3 simulations were run per modification.

The length and amino acid composition of loop 1 varies widely among myosins. A set of studies on loop 1 sequences (43, 44) demonstrated that the length and composition of loop 1 regulate ADP release: shorter loops were associated with decreased ATPase activity, sliding velocity in *in vitro* motility assays, and preferential binding of ADP relative to ATP. Therefore, we anticipate that even low occupancy of these sites has the ability to fine tune contractile performance by modifying ATP cycling.

#### Lysine 951

Lys 951 is located in S2 within the myosin tail (Figure 4). In mature myosin the tail forms a coiled-coil structure where the helical tails of two myosin heavy chains are wrapped around one another. Archetypal coiled-coil helices have a conserved sequence repeat of 7 amino acids: positions *a-g* in which positions *a* and *d* are hydrophobic residues that form a ‘knobs in holes’ interlocking structure that promotes a well-packed hydrophobic core (45). Myosin tails generally follow this archetypal pattern, but there are a small number of ‘skip’ residues that intermittently disrupt the heptad repeat pattern and increase local flexibility into the tail by weakening interactions in the hydrophobic core of the thick filament (46). In implicit solvent MD, the simulated S2 fragment was flexible (Table 4), larger magnitude C_α_ RMSDs were associated with bending of the tail. Although bending occurred, the coiled-coil structure was largely preserved in the unmodified simulations (Figure 9A). Some coiled-coil structure was lost in the K951-Ac simulations in the vicity of the modification site (Figure 9B). In both modified and unmodified simulations there was partial unfolding at the C-terminal end of the fragment. This is likely due to the truncation of the structure or the exclusion of the crystallographically present Hg atoms. In simulations withut PTMs, K951 formed transient interactions with E949 of the opposite chain, but most interactions made by K951 were hydrophobic and involved L950 of the opposite chain (Figure 9A and C). Acetylation of K951 had three effects on the structure of the tail. First, contacts made by K951 were altered: the frequency of salt bridge formation with Glu residues was diminished and K951-Ac interacted more frequently with other residues in the same chain as opposed to the opposite chain (Figure 9B and C). Second, the C-terminal end of the helix became bent and deviated from the typical coiled coil structure. Third, the interhelical distance (measured by calculating the distance between C*_α_* pairs in the two helices) increased in the presence of the PTM (Figure 9D and E). Additionally, K951-Ac altered the electrostatic potential of the S2 fragment, and notably the affected region is proximal to the interacting myosin heads in the super relaxed conformation (Figure 9F). K951-Ac subverted the local coiled-coil structure of S2.

**Figure 9.**
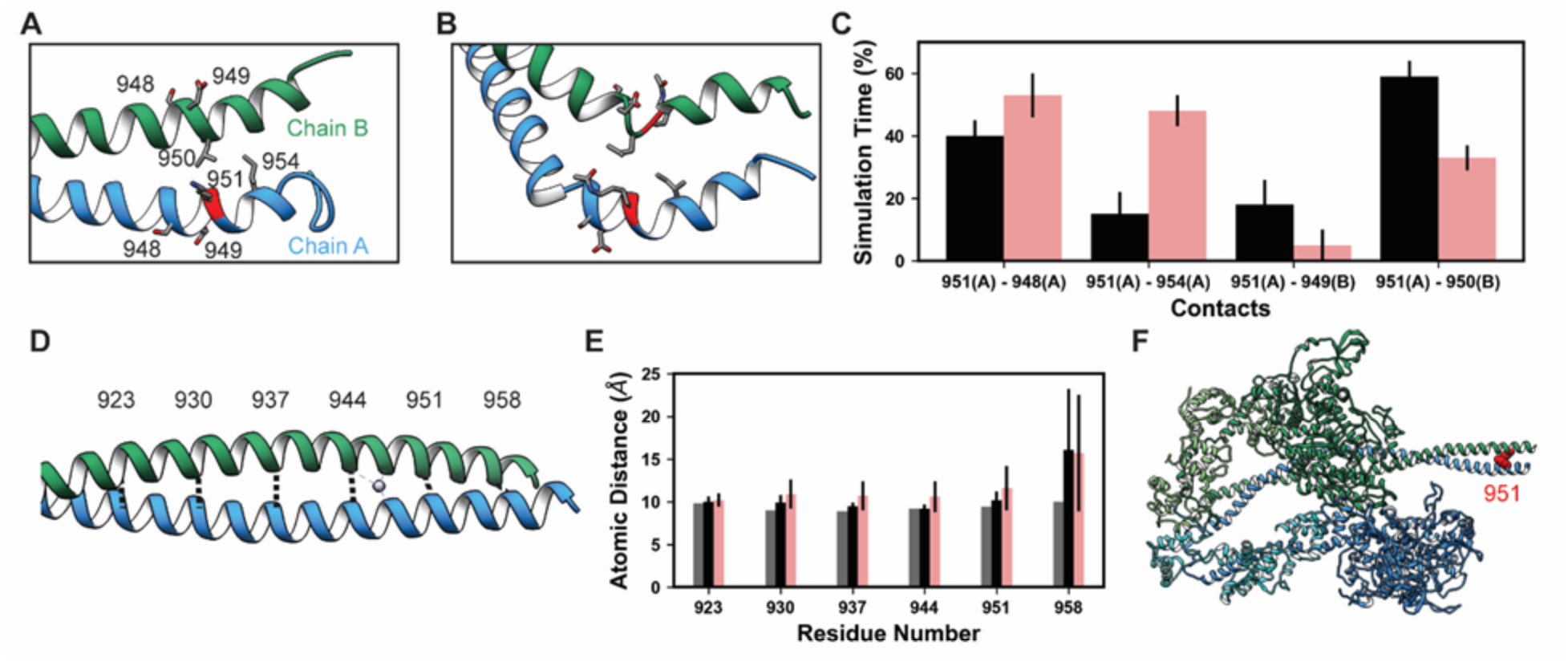
K951-Ac increased flexibility of the myosin tail. Structural changes in the S2 fragment caused by K951-Ac are shown in representative shapshots from MD simulations. In the snapshots, the ribbon of residue 951 is colored red, chain A is colored blue, and chain B green. The atoms of neighboring sidechains are displayed. The coiled-coil structure was preserved in the unmodified simulations (A). In the modified simulations (B), the coiled-coil structure was interrupted by kinks, a loss of *α*-helix structure, and increased separation of the chains. Collectively, these changes increased the local flexibility of S2. (C) Changes in S2structure and dynamics caused by K951- Ac were associagted with changes in local inter-and intra-chain contacts. (D) The inter-chain distance was monitored at 6 positions along S2, chosen to align with the heptad repeat position of the modified residue position. Distances between the C*α* atoms between these residues were tracked for the X-ray structure (shown) and the MD simulations. (E) K951-Ac increased the inter- helix distance (∼1Å) relative to the unmodified simulations. This effect propagated towards the N- terminal end of the helix. The effect may also propagate towards the C-terminus; however the structure is truncated at residue 961 and unfolding of the helices occurred in these simulations. (F) In the super relaxed conformation, the portion of S2 affected by this modification is located nearby the motor domains as indicated on this model (PDB ID: 5TBY). Statistical analysis was performed using Student’s t-test and n=3 simulations were run per modification.

## DISCUSSION

Our study shows that the identified phosphorylation and acetylation PTMs are present on β-MHC in non-diseased human hearts in low abundance. The presence of PTMs on β-MHC may fine tune its function, altering the dynamics of some myosin motors in the cardiac sarcomere. Cumulatively, the functional alteration of a select number of *β*-MHC molecules may be a “direct readout” of the cardiac cellular environment that dictates the needed changes in PTM addition/removal in context with degree of pathological insult and remodeling. Therefore, the normally low levels of PTMs on cardiac *β*-MHC suggest they may contribute to the normal functional and contractile state of the protein and muscle filaments that contain them. Our modeling and molecular simulations provide valuable insight on the functional changes that these PTMs might impart. In our study, the majority of the PTMs on *β*-MHC were reduced with cardiac disease development, which may correspond with increasing expression of *α*-MHC (*MYH6*) in the failing human heart. Therefore, it remains unknown whether reduced PTM abundance at these sites accompanied and/or precipitated the functional decline in the non-ischemic and ischemic failing hearts or rather was a secondary result of the altered cellular environment.

### Extrapolation of Function – Comparing Known Cardiomyopathy-Associated Variants with Identified PTM Sites

To gain further insight on the potential significance of the PTMs identified on cardiac β-MHC, we compared our results with nearby known variants linked to cardiomyopathic diseases in humans. There are numerous pathogenic variants in human cardiac slow β-MHC, the predominant isoform in the myocardial ventricles. However, the study of the function of human cardiac β-MHC has been challenging since most *in vivo* studies of these variants have been performed in mice, which predominantly express the faster isoform α-cardiac MHC (47). This has posed issues when interpreting the effects of β-cardiac MHC variants on cardiac function. *MYH7* variants are implicated in roughly one-third of diagnosed familial HCM cases (48) and 10% of familial DCM cases (49). Ventricular samples obtained during cardiac surgeries in cardiomyopathy patients have been utilized to establish how pathogenic variants alter structure-function relationships that lead to impaired cardiac muscle function and hypertrophy (5). In this study, we compared the impact of cardiomyopathic variants located in the same region as the PTMs we identified, e.g. DCM-linked variants T412N in the head region and R1193S in the tail domain (50). These variants may impact actin-myosin interactions or impair myosin rod structure and assembly (50). Additionally, we show that the PTMs K58-Ac, S210-P/T215-P, and K429-Ac are located in close proximity to pathogenic variants and are present under non-diseased conditions, and tend to decrease in heart failure in most cases with a significant decrease in K951-Ac. See Supplemental Table 2 for comparisons between location of PTMs found in this study with existing reports of pathogenic variants in the *MYH7* gene.

### Phosphorylation of β-MHC

The earliest reports of myosin phosphorylation occurred in the rod region of molluscan smooth muscles that controlled a specialized stretch-resistant “catch” state capable of maintaining tension for long periods (51, 52). In acanthamoeba myosin II, phosphorylation of a short non-helical C-terminal tail piece of each heavy chain increased the compactness of the protein (53). Additional studies in mammalian systems have indicated that the force/power component of muscle depends upon the positioning of the myosin head (54). Kawai et al. utilized liquid chromatography-tandem mass spectrometry and found that the rod region of α-MHC was hypophosphorylated in the HCM-linked cTnC-A8V mouse model compared to controls, the predominant myosin isoform in the adult mouse heart. Left ventricular papillary muscle bundles used in the phosphate study exhibited perturbed cross-bridge kinetics (21).

Several studies have provided insight on how phosphorylation of cardiac proteins alters stretch activation during twitch-contractions in intact cardiac muscle. The Frank-Starling mechanism may in turn be modified to increase cardiac output under conditions of increased venous return (55). Incidentally, phosphorylation of myofilament proteins by PKCβII and PKA influences length-dependent prolongation of heart muscle relaxation (56, 57). Sarcomeric protein phosphorylation is regulated by reactive oxygen species (ROS), whereby oxidative stress tends to increase protein phosphorylation due to inhibition of protein phosphatase activation and stimulation of protein kinases (7). Crosstalk between PTMs may in fact provide a higher order of regulation, e.g. same-site competition, structural changes in secondary sites that make it more accessible for modification by other PTMs, or direct modification of the modification of a secondary PTM (58).

### Acetylation of β-MHC

In vitro studies have identified potential modification sites that include glutathionylation and glutathione of residues in the myosin head. However, these findings have yet to be verified *in vivo* (59). Lysine-acetylation/deacetylation contributes to cardioprotection and cell survival during ischemic insult. Recent reports demonstrated increased levels of lysine acetylation (acetyl-Lys)-dependent activation and AMP:ATP that activates a network that includes AMP-activated protein kinase (AMPK), AKT and PKA kinases during ischemia (60). Moreover, there is evidence of cross-talk between phosphorylation and acetyl-Lys that can affect recruitment of kinases/phosphatases by acetyl-Lys (61, 62) or recruitment of HDAC/HATs by phosphorylation (63). Studies using MALDI-TOF/TOF MS have detected an increased number of acetylated proteins in two distinct rodent models of induced pressure-overload cardiac remodeling that may be linked to metabolic changes associated with heart failure (64). These alterations after pressure-overload may be due to reduced abundance of the mitochondrial deacetylase, NAD-dependent deacetylase sirtuin-3 (Sirt3). These studies, however, did not examine proteins with reduced protein-acetylation upon ischemic or non-ischemic cardiomyopathy-induced heart failure (64).

The levels of acetylated proteins in the heart examined in this study depends on HDAC activity as much as HAT activity. Class I HDACs promote pathological cardiac hypertrophy, while Class IIa HDACs suppress cardiac hypertrophy (62). Studies examining the effect of sarcomere protein acetylation and class I and II HDAC inhibitors on cardiac function concluded that this treatment increased Ca^2+^-sensitivity of isolated myofilaments (9). The same group found the Class II HDAC4 associated with the sarcomere (8, 9). Recent studies have explored the functional significance of sarcomeric protein acetylation to understand the impact of histone deacetylases (HDAC) on cardiac function (65). Administration of the histone deacetylase inhibitor ITF2357 (Givinostat) improved heart relaxation in heart failure in rodent models with preserved ejection fraction (HFpEF) by promoting myofibril relaxation (65). HDAC inhibitors are beneficial to cardiac function by targeting cytosolic and sarcomeric proteins by improving muscle contractility (10) and relaxation (65) as well as cardio protection during ischemia/reperfusion injury by promoting autophagy (66). Therefore, it is reasonable to suggest that a reduction in acetylation at K951 and K1195 may be associated with declining myocardial function. β-MHC may be included in the list of sarcomeric proteins that benefit from HDAC-inhibition. In our study we see distinct differences in β-MHC acetylation that seem to be influenced more by location of the lysine than the the type of heart disease (ischemic- and non-ischemic-HF). In control donor hearts the residues K951 and K1195 in the myosin rod region are acetylated in ∼14% and 2% of the peptides, respectively. The rod region is perhaps the more conformationally consistent portion of the myosin protein that is accessible to HATs but also HDACs. Therefore, one interpretation of the tendency for lower acetylation of these residues in the failing heart samples is that they are not in the globular domain and may be more accessible to HDACs. It is unknown whether HDAC-inhibition would preserve acetylation at these sites, and whether maintenance of these PTMs is functionally important.

K58 acetylation on the other hand is markedly reduced in the ischemic-HF hearts, and its localization near the SH3 domain may also be slightly more exposed, with potentially greater HDAC activity in ischemic hearts. The other acetylated residues K34 and K429 are located further into the interior of the myosin head and may be less affected by increased HDAC activity in diseased hearts. Preservation of acetylated lysines in *β*-MHC may be beneficial to normal heart function and pathological conditions favoring greater HDAC activation may disrupt the normally low levels of acetylated residues decreasing Ca^2+^ sensitivity and impacting relaxation in heart failure.

### Molecular Dynamics Simulations to Describe the Functional Impact of β-MHC PTMs

Molecular dynamics simulations were performed to better understand the functional significance of the β- MHC PTMs. Focus was placed on modifications in the regions with greatest functional significance (K213-Ac and T215-P at ATP-binding pocket) and K58-Ac (SH3 domain in the head region) as well as modified residues with significantly altered abundance in diseased states (K951- Ac). At the ends of each S1 head is the lever arm domain composed of an α-helical coiled-coil tail. Movement of the lever arm transduces energy garnered from ATP hydrolysis into motion and allows production of force. The stability of the lever arm is governed by the two light chains named the ELC and the RLC that bind to the lever arm domain in tandem (67), which also dictate the positions of the myosin head relative to the myosin rod and actin filament (13). Because the LCs influence myosin head positioning they can modulate speed and contractile force of cardiac muscle by turning contraction “on” or “off” in response to changes in intracellular calcium levels.

Lowey *et al*. and others have provided evidence using cryo-electron microscopy and variant isoforms of the ELC that SH3 domains of class II myosins interact with an N-terminal extension present in a cardiac ELC isoform (68, 69). They have proposed that an N-terminal sequence of the ELC (^4^`KKPEPKK` in human cardiac and ^4^`KKDVKK` in chicken skeletal muscle) transiently interacts with actin filaments and modifies the kinetics of the chemo-mechanical cycle. They posit that interaction between the ELC and actin is mediated by the SH3 domain of myosin S1: specifically, the SH3 domain binds to a poly-Pro rich sequence in the N-terminus of the ELC and optimally positions the N-terminus of the ELC for interaction with actin. The ELC does contain several sequence motifs that correspond to SH3 targets, including ^21^`PAPAPP` and ^26^`PPEPERP` (potential motifs were identified using the *Linear Motif Domain Interaction Prediction* algorithm).

The K58-Ac simulations performed using the X-ray crystal structures of the β-MHC indicated that acetylation of Lys 58 caused minimal structural perturbations overall compared to the non-modified protein. Neutralization of K58 charge and introduction of the acetyl group appeared to reduce flexibility of adjacent residues C79-82, but it did not appear to significantly perturb motor domain conformation and dynamics. Overall, the greatest impact of K58Ac is the decrease in electrostatic potential and structure on the outer surface of the SH3 domain, which is expected to alter interactions with sarcomeric regulatory proteins it may contact. *In vitro* studies from Lowey *et al.* showed that interactions between the ELC and F-actin slowed the shortening velocity in skinned muscle fibers and that the interaction reduced velocity in *in vitro* motility assays (68). In light of this, our simulations suggest that acetylation of K58 has the potential to impede interactions between the ELC and SH3-like domains, thereby reducing interaction between the ELC N-terminus of actin filaments and effectively increasing shortening velocity. This prediction, however, is putative, as we have not tested the affinity of SH3 variants for potential ELC recognition sequences. This prediction is based off of the observed covering of the putative binding pocket by K58-Ac. One of the *LMDIP-*predicted SH3 binding sequences includes two Glu residues, and we can predict that K58-Ac would also have reduced affinity for this peptide due to the loss of the charge on Lys. Charged residues can play roles in restricting the specificity of SH3- like domains. Because we have not directly simulated binding between SH3 and the ELC, it remains possible that acetylation could increase affinity of SH3 for the ELC via altered intermolecular interactions, but we find this the less likely scenario. Future computational studies investigating K58-Ac should examine binding between SH3 and potential recognition sequences in the ELC. Similarly, future *in vitro* studies could confirm or deny our computational predictions by measuring sliding velocity in *in vitro* motility assays with modified and unmodified K58 residues. We suggest that K58-Ac may provide a reversible means to decrease the electrostatic potential of the SH3 domain surface, which may alter its interactions/affinity for ELC and perhaps other sarcomeric regulatory proteins.

The doubly modified peptide K213-Ac/T215-P was deemed to contain high-confidence PTM sites, however S210 and T215 were found detectable but not quantifiable. Given the high functional significance of their location, loop 1 within the ATP binding pocket even low abundance modifications are expected to exert a palpable effect on the myocardium and tune its contractile responsiveness. In our simulations, the modified loop 1 formed a more compact structure and we predict that these PTMs would effectively behave as shorter loops that stabilize ADP. However, our simulations were performed in a post-rigor-like ATP-bound, actin-free structure of myosin and have insufficient sampling to measure ATP affinity or ATP binding/ADP release rates. Additionally, the rules that govern the relationship between loop 1 and nucleotide binding have not been definitely established: it is not immediately clear what interactions made by loop 1 specifically regulate ADP release, which complicates *in silico* predictions made here. Future studies on these post-translational modifications in loop 1 should investigate their effects on ATPase activity and ADP release rates. We speculate that these structural perturbations in the ATP binding pocket could alter the ADP-release rate. Decreases in the ADP-release rate could prolong the power stroke and increase force, whereas increases in the ADP-release rate could diminish force generating capacity of the muscle. It remains unknown how low levels of phosphorylation at S210 would affect the structural dynamics of the ATP binding pocket region if S210-P, K213- Ac and T215-P were simultaneously modified. While in our sample set we do not observe any significant change in amount of T215-P in the different diseased states the variability of abundance in non-failing controls prevented drawing any valid conclusions. It certainly could be predicted that if low level modification of these residues could transiently fine tune or tweak force production by β-MHC it could be advantageous, a reduction in the diseased states would therefore suggest the loss of one out of potentially many mechanisms that modulate β-MHC function.

Our simulations suggest that acetylation of K951 disrupts the native structure of the coiled coil and increases flexibility within the tail. Increased flexibility within the tail may alter the orientation of myosin heads relative to the thin filament, alter association rates between the thick and thin filament, and may affect the ability of myosin to form the interacting heads motif. Our simulations alone are not able to assess which of these effects on myosin dynamics is most likely due to the limited fragment of myosin that was simulated. Limitations of these simulations are as follows. First, we have only simulated a fragment of the tail. Crystal contacts and the Hg atoms in the x-ray structure may have contributed to the stability of the tail. We did observe transient unfolding at both the N- and C-termini in our simulations. Unfortunately, K951 is located close to the C-terminus of the simulated structure and some of our observations may be attributed to the effects of tail truncation. Next, in our simulations both K951 in chain A and B were modified, so our simulations sample the most aggressive effects of acetylation at this site: diminished effects may be found for singly modified systems and there may be asymmetric behavior depending on which chain is modified. Finally, because our simulations only incorporate a fragment of the tail, it is difficult to predict the effects of reduced packing on overall tail flexibility, the orientation of myosin heads, or effects on access to the interacting heads motif. Nevertheless, our simulations do provide strong evidence for increased flexibility in the tail. We attribute this to altered electrostatic interactions in the vicinity of K951. Acetylation of K1195 may result in similar effects. This residue is nearby skip 1 of the myosin rod. Post-translational modification of this region may further modulate tail flexibility or the assembly of the myosin rod (46).

### Conclusions

Our study identified novel PTMs on *β*-MHC in non-failing and failing human hearts. Addition of PTMs to β-MHC may be an energetically conservative approach to modify myosin motor function that potentially precedes switching to the fetal isoform α-MHC in the failing heart. Overall, there tended to be less PTM abundance in the heart failure conditions examined here, which may be due to increased phosphatase or HDAC activity. Reduction of these PTMs in heart failure may be compensated for by the documented increase of α-MHC expression that occurs in human heart failure. Our modeling data suggest that some of these PTMS have potential to alter dynamics within β-MHC and thereby fine tune myofilament function. It remains to be seen whether loss of these PTMs in failing hearts represents the removal of a beneficial, albeit subtle regulation of myofilament function.

## Acknowledgements

The authors wish to thank the Lifeline of Ohio for the collaboration on non-failing donor tissue, and surgeons and transplant coordinators at the Ohio State University Wexner Medical Center for helping obtaining the end-stage failing tissue.

## Abbreviations

Aa: amino acid
Ac: acetylation
AMPK: AMP-activated protein kinase
β-MHC: beta myosin heavy chain protein
CID: collisional-induced dissociation
cTn: cardiac troponin complex
cTnC: cardiac troponin
C cTnI: cardiac troponin I
cTnT: cardiac troponin T
cTnT3: human cardiac troponin T isoform 3
cTnT4: human cardiac troponin T isoform 4
DCM: dilated cardiomyopathy
ELC: essential light chain
HAT: histone acetyltransferase
HCM: hypertrophic cardiomyopathy
HDAC4: histone deacetylase 4
HDAC6: histone deacetylase 6
HF: heart failure
HFpEF: preserved ejection fraction
IRP: internal reference peptide
I-HF: ischemic heart failure
IRC: internal reference control
LC: light chain
LC/MS: liquid chromatography-mass spectrometry
MALDI-TOF/TOF MS: MALDI coupled to time-of-flight mass spectrometry
MD: molecular dynamics
MHC: myosin heavy chain
MI: myocardial infarction
MLCK: myosin light-chain kinase
MS: mass spectroscopy
MyBP-C: myosin-binding protein-C
MYH7: beta myosin heavy chain gene
Myosin-S1: myosin subfragment-1
Myosin-S2: myosin subfragment-2
NF: non-failure
NI-HF: non-ischemic heart failure
P: phosphorylation
PCAF: p300/CBP-associated factor
PKA: protein kinase A
PKCβII: protein kinase C beta II isoform
PTM: post-translational modifications
RLC: regulatory light chain
RMSD: root-mean-squared deviation
RMSF: root-mean-squared fluctuation
ROS: reactive oxygen species
SASA: solvent accessible surface area
SEC: size-exclusion chromatography
Sirt3: sirtuin-3
TCA: tricarboxylic acid cycle
Tm: tropomyosin
WT: wild type

---

**Supplemental Figure 1.**
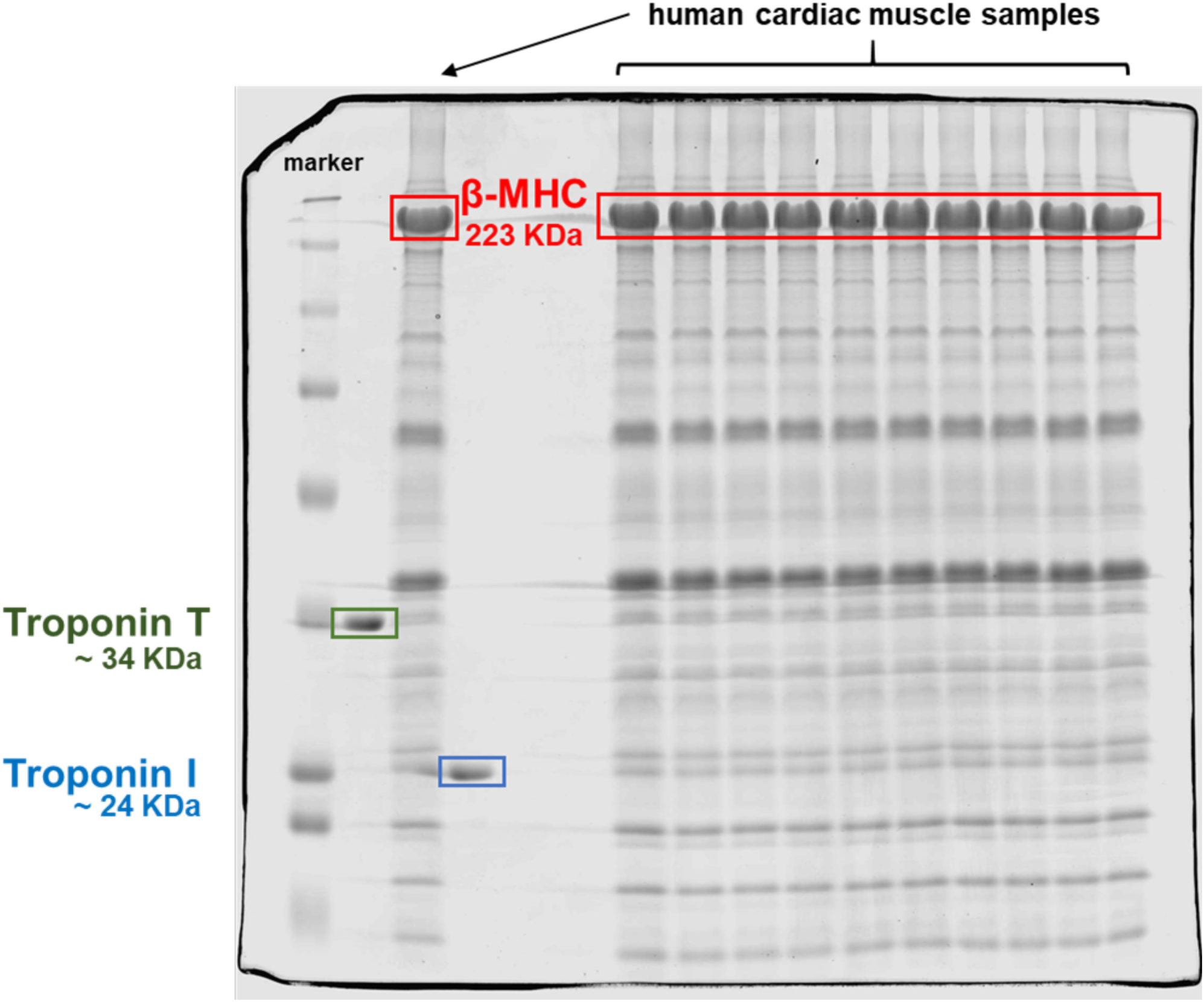
Coomassie-stained gel of homogenized human heart tissues. The red box indicates the band corresponding to β-MHC in the cardiac human samples analyzed here at the predicted molecular weight of ∼ 223 kDa. The lanes are: BioRad Dual marker, TnT (green box, ∼34 KDa), cardiac human sample, TnI (blue box, ∼24 KDa), two blank lanes, then ten lanes of human cardiac samples.

**Supplemental Figure 2.**
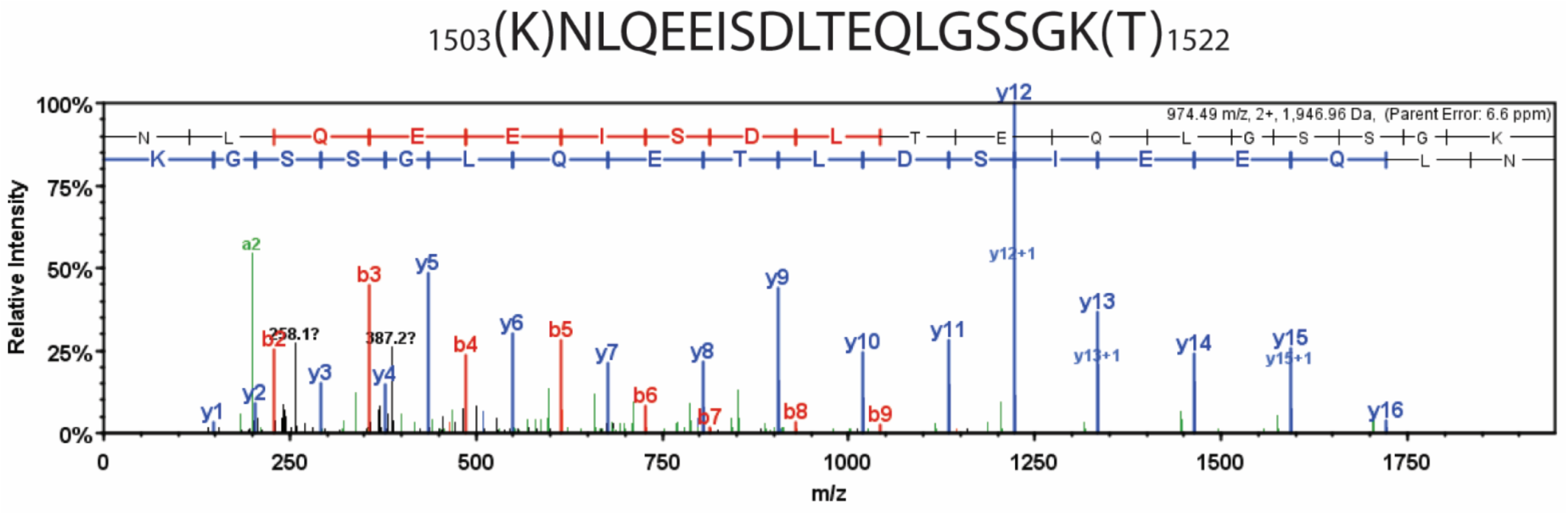
Detailed liquid chromatography-mass spectrometry spectrum of the common internal peptide sequence (IRP). MS/MS spectrum of trypsin-digested β-MHC peptide sequence (1504-1521, m/z 974.49). b-ions and y-ions indicate N-terminal and C-terminal ions, respectively.

**Supplemental Figure 3.**
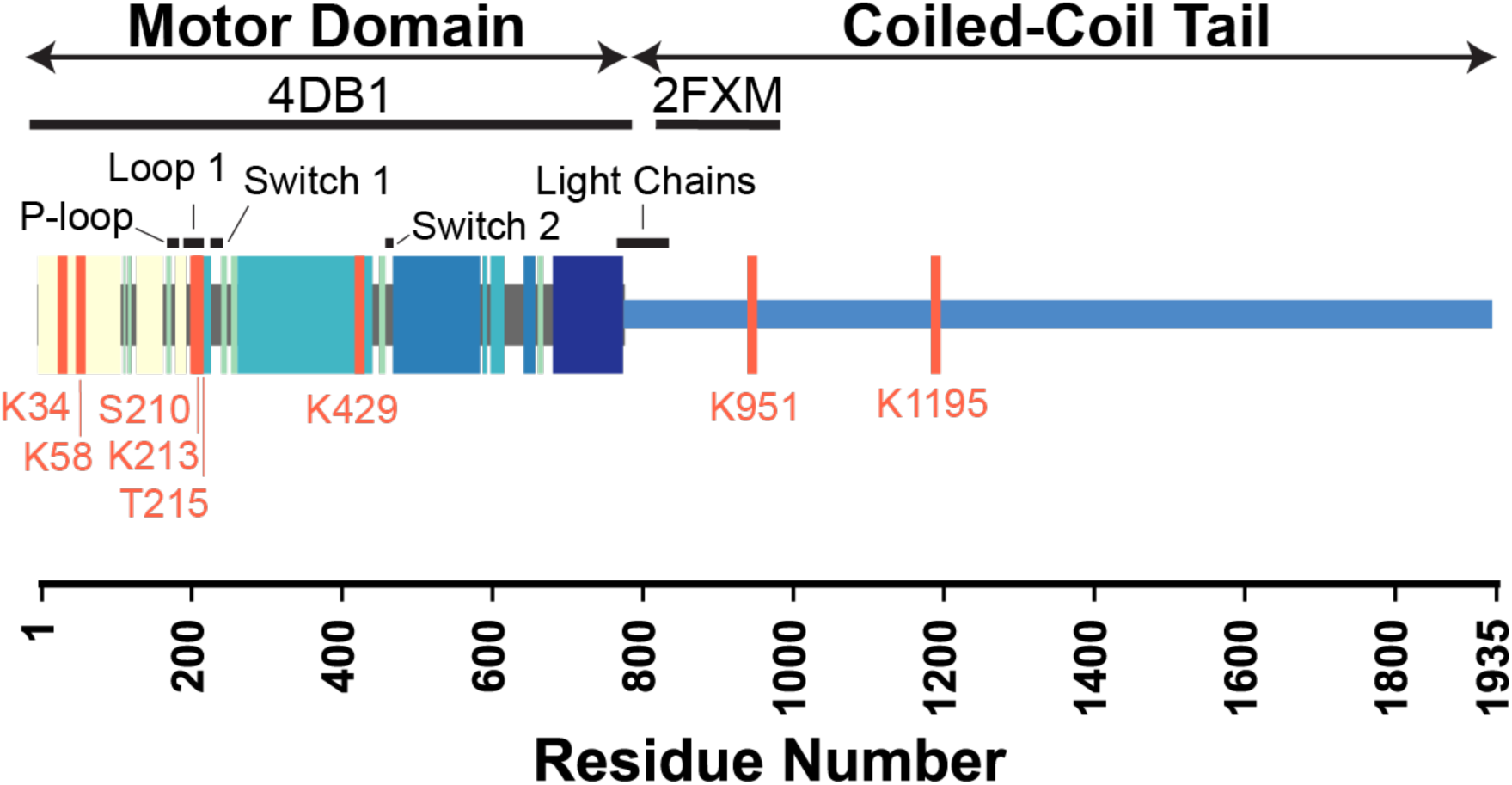
Key functional regions of cardiac β-myosin motor domain and locations of the identified PTMs. The locations of PTMs and functional regions are annotated on a schematic of the *β*-MHC sequence. PTM sites are denoted with pink lines and significant structural elements are annotated with black bars. The regions of the sequence modeled in our simulations (4DB1 and 2FXM) are denoted with thick black bars. Colored blocks correspond to structural elements shown in Figure 4 panels A and B.

**Supplemental Figure 4.**
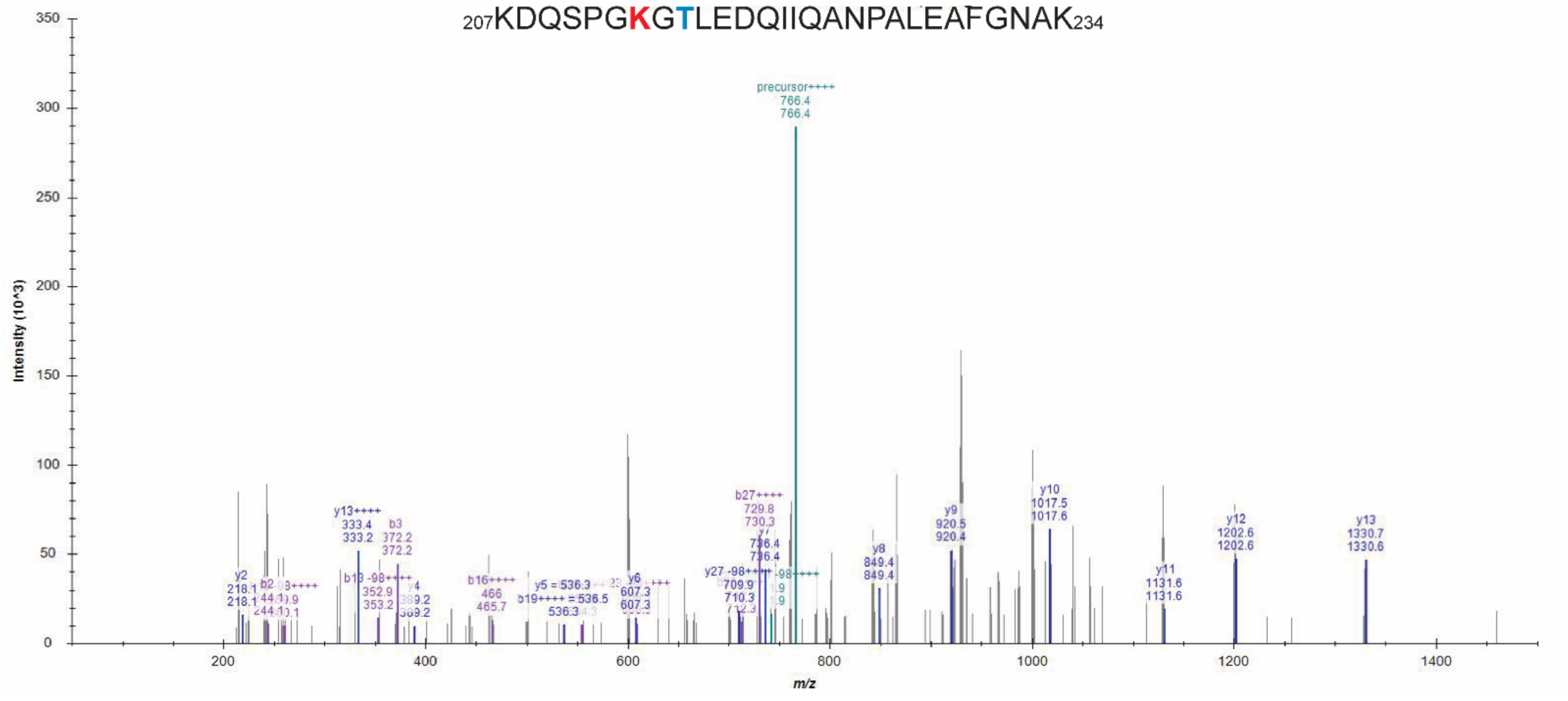
MS/MS spectrum of the doubly-modified peptide sequence. The K213-Ac/T215-P modified sequence (207-234, m/z 766.36) presented an isotope dot product (idotp) value below 0.5 which indicates that the modified peptide was detectable but not quantifiable. Acetylated residues are highlighted in red while phosphorylated residues are found in blue.

**Supplemental Table 1.** Summary of the patients’ demographic features. De-identified human heart samples were obtained from non-failing, ischemic heart failure, and non-ischemic heart failure patients.

| **Sample #** | **Failing Category** | **Age** | **Gender** | **Race** |
| --- | --- | --- | --- | --- |
| **1** | Non-failing | 51 | F | White |
| **2** | Non-failing | 69 | M | White |
| **3** | Non-failing | 41 | M | Black |
| **4** | Non-failing | 58 | M | White |
| **5** | Ischemic heart failure | 50 | F | White |
| **6** | Ischemic heart failure | 68 | M | White |
| **7** | Ischemic heart failure | 47 | M | Black |
| **8** | Ischemic heart failure | 62 | M | Black |
| **9** | Non-ischemic heart failure | 53 | F | Black |
| **10** | Non-ischemic heart failure | 68 | M | White |
| **11** | Non-ischemic heart failure | 47 | M | White |
| **12** | Non-ischemic heart failure | 61 | M | Black |

**Supplemental Table 2.**
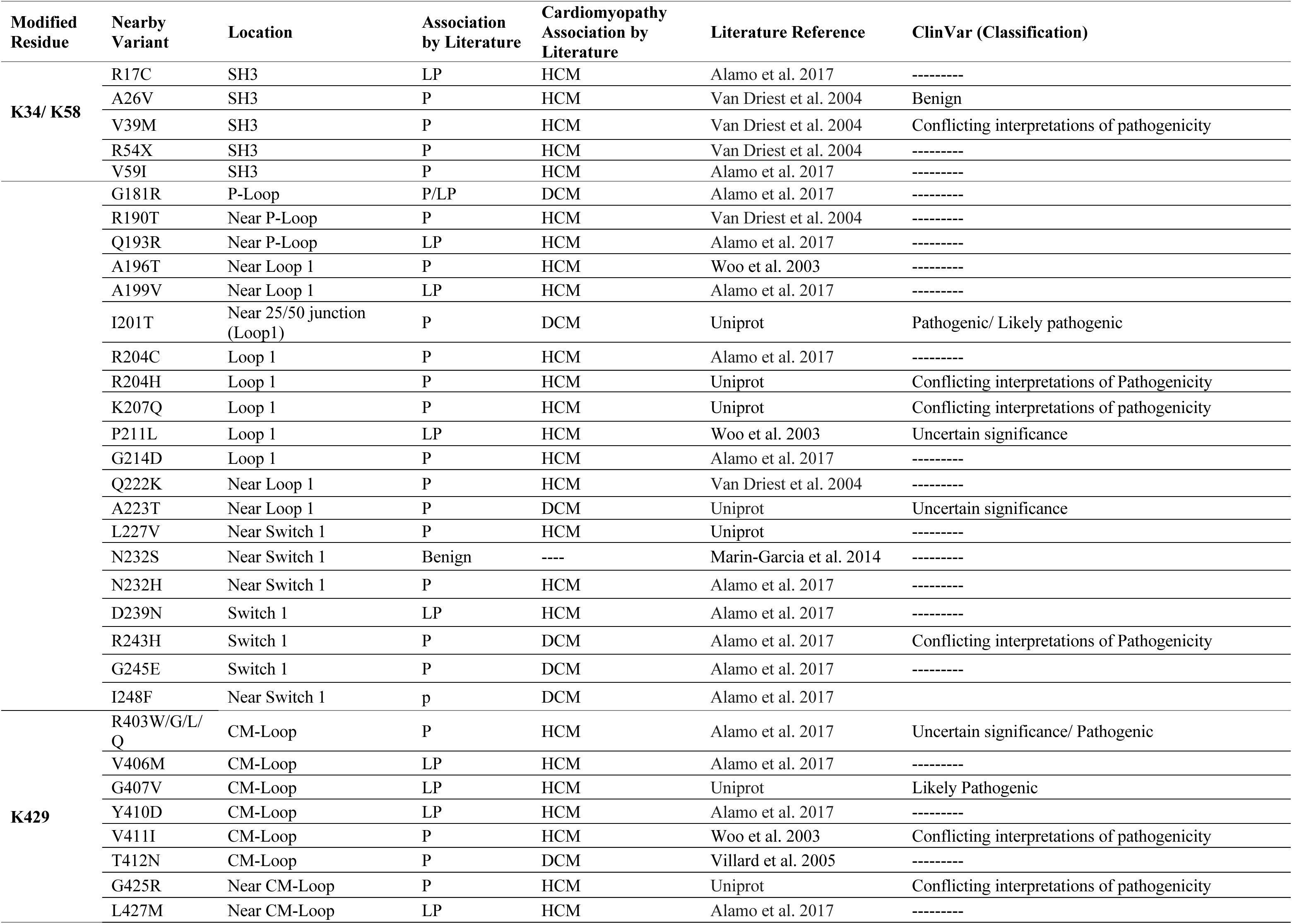
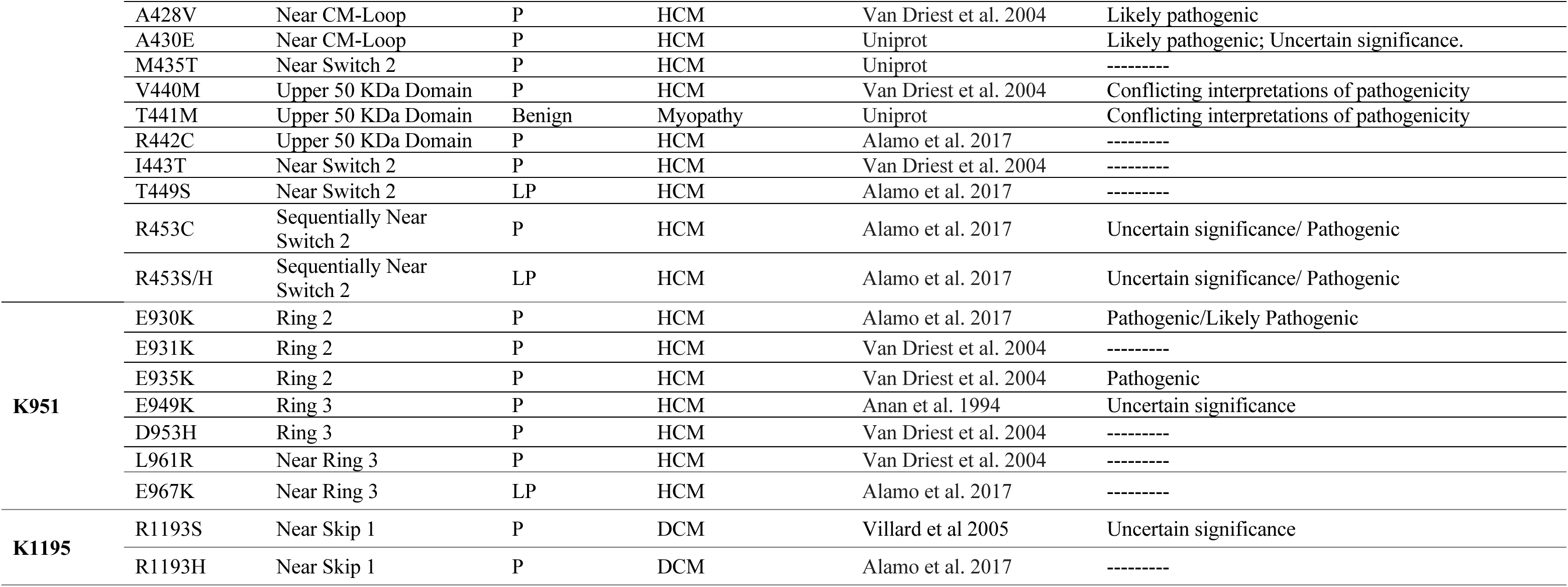
Comparison of location of residues bearing PTMs with known cardiomyopathy variants in *MHY7*. List of potential pathogenicity of the variants and their location within nearby PTM regions. Abbreviations are the following cardiomyopathy-loop (CM-loop), likely pathogenic (LP), pathogenic (P), hypertrophic cardiomyopathy (HCM), and dilated cardiomyopathy (DCM).

